# Multi-objective optimization-based design of a compliant gravity balancing orthosis: development and validation

**DOI:** 10.64898/2026.03.19.712706

**Authors:** Haider A. Chishty, Zion D. Lee, Uday Kiran Balaga, Fabrizio Sergi

## Abstract

Wearable devices for gravity balancing have high potential for impact across domains, including neuromotor rehabilitation and occupational systems. Devices made from compliant mechanisms, optimized to achieve specific compensation moments at target joints, have proven effective, but thus far have solely been optimized towards gravity compensation and not other wearability criteria. In this work, we propose a multi-objective optimization framework, based on particle swarm optimization, to design a soft, gravity balancing shoulder orthosis, while taking into account wearability constraints such as undesired loading directions and device size. Using this custom framework, we pursued multiple stages of orthosis design and optimization, selecting multiple solutions to be translated to real-world prototypes. These solutions were realized via 3D printing with thermoplastic polyurethane and evaluated for mechanical performance on benchtop and in-vivo. In-vivo testing on 6 healthy individuals demonstrated relative reductions in muscle activity for the anterior deltoid and upper trapezius, by 53 % and 71 % respectively when operating the orthosis for static tasks within functional shoulder ranges of motion. Changes in muscle activation were also were observed across other muscles, including the posterior deltoid, as well as in dynamic tasks at different speeds.

## I. Introduction

Shoulder disability and weakness in the surrounding muscles can severely inhibit one’s ability to perform necessary activities of daily living (ADLs) [1]–[4]. Such conditions can be due to orthopedic or neurological disorders; the latter can result in motor function deficits, such as those caused by stroke, which can lead to weakness on one side of the body or abnormal synergies between muscle groups [4]–[7].

In cases of shoulder disability, maintaining arm elevation is difficult [4]. Assistive devices that provide restorative forces and moments to the shoulder have been shown to facilitate functional arm movement, reduce muscle effort, and restore ADL performance [1]–[4]. Such support has been applied in rehabilitation, where it helps decouple stroke-related muscle synergies [5], [7], [8], and in occupational settings, where shoulder exoskeletons reduce musculoskeletal injury risk [2], [9]–[11].

Occupational exoskeletons typically rely on passive elastic elements [11]. In wearable support design, avoiding the complexity and bulk of active systems is advantageous [3]. Passive elastic mechanisms are increasingly used for shoulder support due to their compactness [12], though achieving the desired nonlinear compensation requires careful geometric design. Elastic components may be mounted directly on the arm [12], [13], as implemented in devices such as the WREX, Armeo Spring, and ShoulderX [9], [14], [15]. Alternatively, components can be positioned proximally and routed through variable-radius pulleys to create modular moment arms, as in the WPCSE, AD Exo, and HIT-POSE [4], [10], [11].

An alternative gravity-compensation strategy leverages compliant mechanisms, which exploit structural flexibility and nonlinearity to avoid the large rigid elements common in spring-based designs [1]–[3], [16]–[19]. These mechanisms store elastic energy through large deformations while transmitting motion and force [17], [20], offering advantages in wear, friction, reliability, and manufacturability [17], [20]. Their force–displacement behavior has been extensively studied, enabling their use as spring alternatives in statically balanced systems [19], [20] and in mechanisms that generate prescribed load paths [18], [19]. These principles have been translated to in-vivo applications, where compliant elements function as torsional joint springs [1]–[3]. Achieving specific output loading requires coupling finite element or similar simulations with optimization frameworks [1]–[3], [17]–[19]. To date, these frameworks have prioritized gravity-balancing performance and have not explicitly focused on wearability constraints such as size or undesirable loading.

We present an optimization-based framework that integrates finite element modeling with multi-objective optimization to design comfortable and practical gravity balancing shoulder orthoses. A sequence of increasingly complex optimizations was performed to identify the simplest configuration capable of producing feasible designs. Finally, a multi-participant validation study evaluated the orthosis’ effects on key muscles involved in shoulder flexion.

## II. Methods

### A. Problem Definition

As the moment needed to support the shoulder against the arm’s weight is dependent on both shoulder and elbow angles, it is useful to first simplify the problem. This was done by designing the framework to balance a 1 degree of freedom (DOF) system; that is, constraining shoulder motion to a single DOF and enforcing the elbow to remain extended. For the purposes of this work, the shoulder DOF that was targeted for support was shoulder flexion.

As seen below, even in the simplified 1 DOF problem, the moment required to balance the system is nonlinear, as it is proportional to the sin of the joint angle (Fig. 1). As such, the moment *M*_*ideal*_ needed to balance the shoulder against the arm’s weight is defined as:

**Fig. 1.**
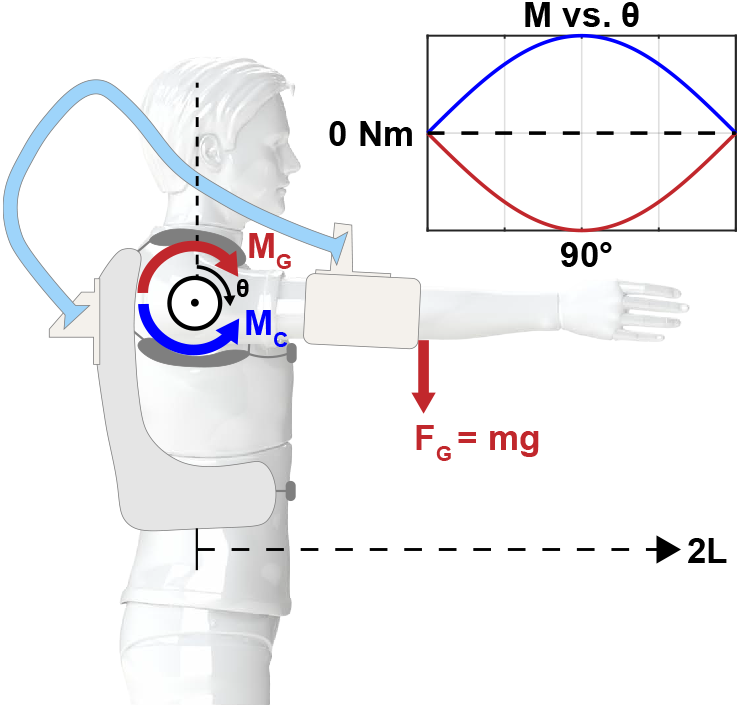
The system to be balanced is simplified to a 1 DOF system. The orthosis (light blue) is a single compliant element that attaches to the user at a proximal (torso) and distal (arm) location. Perfect compensation occurs when the reaction profile is sinusoidal.

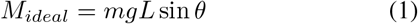

where *m* is the mass of the link, *g* is the gravitational constant, and *L* is half the length of the link. To perfectly support the shoulder, this compensation moment must be implemented in the opposite direction to gravity.

### B. Framework Overview

The framework presented here combines optimization and simulation techniques to identify orthosis designs capable of providing the highly-specific, nonlinear moments required to support the shoulder against gravity while also achieving secondary objectives such as maintaining low spatial profiles and not exhibiting high output loading in undesirable DOFs. The framework integrates particle swarm optimization (PSO) [21] in MATLAB (MathWorks, Natick, MA, United States) with finite element analysis performed in SolidWorks (Dassault Systèmes, Vélizy-Villacoublay, France). Automation and data transfer between software environments was done using a custom-built VB.NET application (Fig. 2). More information on this application is given in Appendix B.

**Fig. 2.**
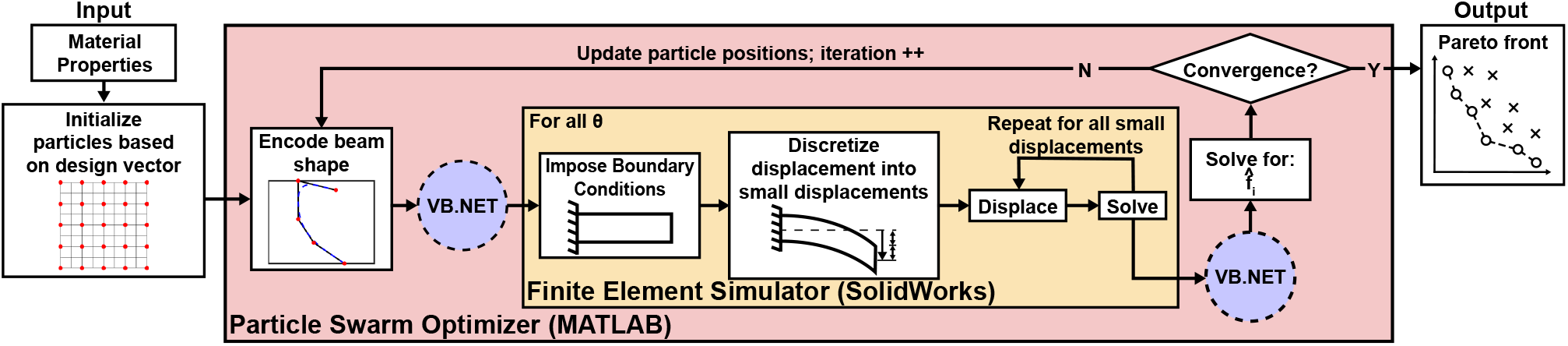
The multi-objective simulation-optimization framework. By implementing a SolidWorks finite element simulation into a particle-swarm optimizer in MATLAB, the framework outputs a Pareto front of orthosis solutions based on multiple design objectives.

The framework was developed in a modular fashion, such that simplification strategies can be enabled or disabled as needed, towards identifying the simplest configuration of process settings required to identify an effective and practical orthosis design. Examples of such strategies include optimization type (Single vs. Multi-Objective), kinematic compatibility (Orthosis Out-of-Plane relative to the arm vs. In-Plane), and specificity of moment amount (adimensional target moments vs dimensional) (Fig. 3).

**Fig. 3.**
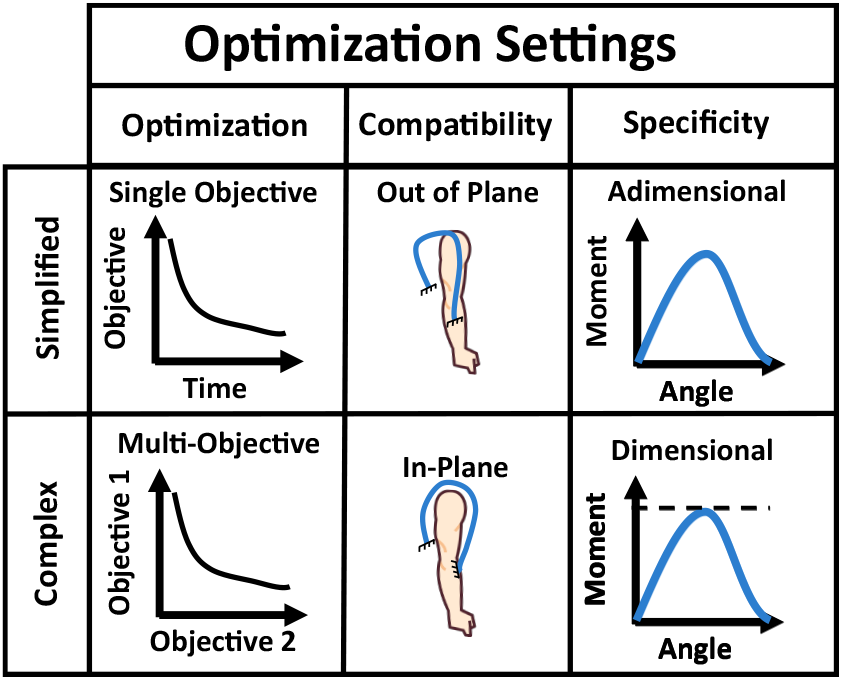
Optimization setting options: for each setting type (optimization, compatibility, and specificity) a simplified and complex version exist.

Towards simplification, this process attempts to leverage the many scaling opportunities available in the 2D problem: for example, for a given orthosis design, scaling the out-of-plane thickness, the in-plane thickness, or the entire design relative to the axis of rotation all scale the resulting compensation moment following simple laws derived from basic solid mechanics principles; however, these scalings may violate size constraints when being used to achieve larger target moments. As such, scaling was initially assumed boundless (in adimensional optimizations), and then later constrained to produce more realistic and directly translatable orthosis solutions (in dimensional optimizations).

The output of the framework is a set of orthosis designs that are optimal in the sense of one or more design objectives. In the case of multi-objective optimization, the output is the Pareto set of the optimization problem; in single-objective cases, sets of solutions classified as differently shaped from eachother are presented. Orthosis designs are classified as differently shaped using a Procrustes analysis - details are given in Appendix A.

### C. Orthosis Shape Encoding

In our framework, we define an orthosis as a monolithic structure, made of an elastic material, which will attach to the user at two ends: 1) the proximal end, attached on the user’s back, and 2) the distal end, attached along the user’s arm (Fig. 1).

The 0° shoulder angle represents the arm completely raised, and requires the minimum amount of compensation moment against gravity; as such this posture was selected to correspond to the undeformed configuration of the orthosis.

Each design is encoded by a design vector (2), with Table I providing descriptions of each component. In general, a linkage chain formulation [18] is used to define the neutral axis of the orthosis. The beginning and end of this control polygon correspond to the orthosis attachment points. The orthosis thickness is allowed to vary using more control points to develop a spline that defines in-plane thickness as a function of orthosis location. Fig 4 demonstrates how the design vector defines an orthosis shape. For this study, *n* = 4 and *m* = 3, yielding a 16-dimensional design vector in the most general case. More information on orthosis encoding, including default design vector bounds, as well as how each component relates to the final orthosis design, is given in Appendix A.

**TABLE I.**
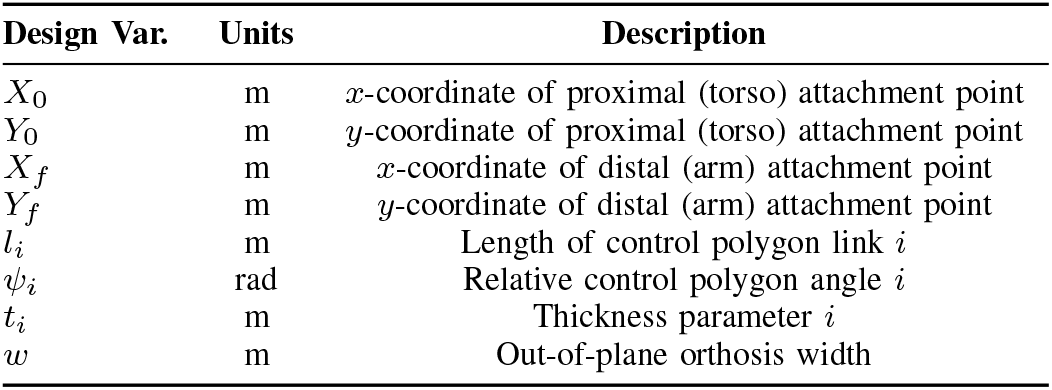
Design Vector Components.

**Fig. 4.**
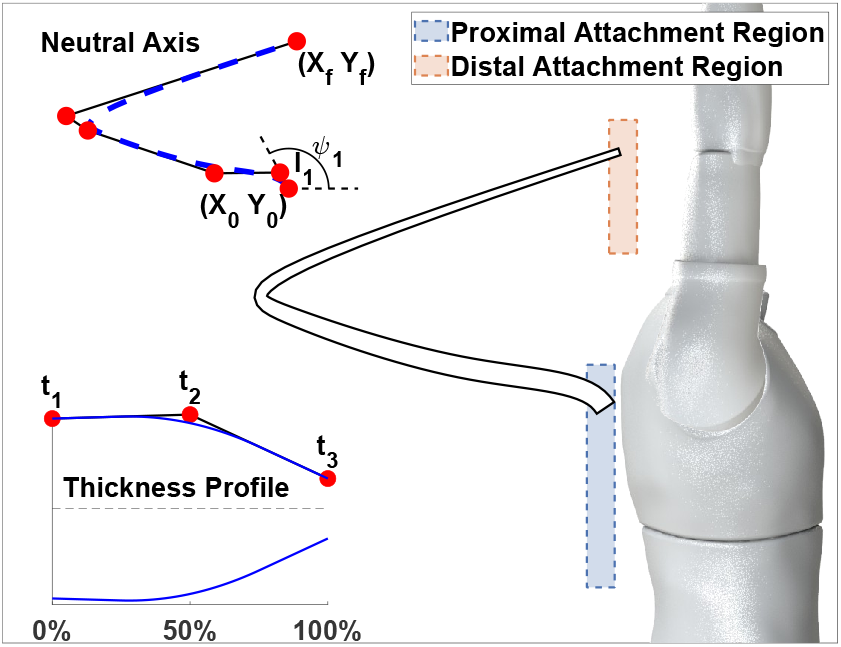
Overall encoding from design vector to orthosis. Orthosis design includes the definition of the neutral axis, defined by basing a b-spline off a linkage chain, and in-plane thickness profile, defining in-plane thickness as a function of neutral axis location, with 0 % and 100 % corresponding to the proximal and distal ends of the orthosis respectively.

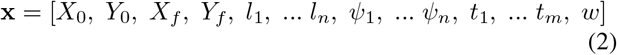

### D. Finite Element Simulations

Finite element simulations were performed in SolidWorks. 3D bodies were subject to 2D, nonlinear, static simulations. Nonlinear simulations apply incremental loading over steps, allowing for a series of deformations to be simulated. This is necessary when an applied deformation is large relative to the size of the body, as is the case for simulations done in this work.

The proximal face of the orthosis is kinematically constrained to not move in any direction. At the distal face, a rotational displacement is enforced, such that the face is rotated 180° clockwise around a remote axis of rotation corresponding to the shoulder. A solid mesh comprised of linear, tetrahedral elements, with a maximum side length (global size) of 15 mm, is created. This configuration was selected as a compromise between simulation time and result accuracy. Material properties corresponding to Flex TPU filament (*E* = 13 MPa, ν = 0.38) from Roboze INC. (Houston, TX, USA) were used for simulation.

Nodal forces from the simulations were used to calculate different metrics of loading, later used used to determine optimization objectives. These calculations are given in Appendix B.

Simulation results were validated on an instrumented bench-top apparatus with small-scale orthosis prototypes (Fig. S6). Experimental methods and results from this validation are given in Appendix C. Observations from these validations were used to tune the material properties used in simulations.

### E. Optimization

#### 1) Objective Calculations

In the following sections we describe the different optimization objectives used in this work, as well as how they are specifically defined.

#### a) Gravity Compensation

The gravity compensation capabilities of an orthosis design are determined by comparing the shoulder compensation moment *M*_*G*_ to the ideal moment profile. This comparison is done as an absolute difference at each simulation step (joint angle increment), normalized by the ideal profile.

For adimensional optimizations, where the ideal profile is a sin wave with a nonspecific amplitude, the cost function for this optimization objective is defined as:

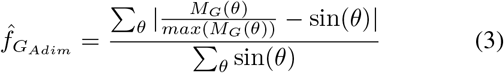

where 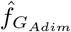 represents the normalized error in gravity compensation accuracy.

For dimensional optimizations, the compensation profile is not normalized by its maximum. Further, an amplitude parameter *A* defining the desired moment compensation amplitude is introduced. For the dimensional optimization used in this work, *A* = 8 Nm, corresponding to the moment needed to accurately compensate the arm of a 50^th^ percentile male [22].

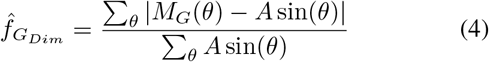

#### b) Concentrated Moment at the Distal End

Based on prototype testing, it was evident that orthoses that exhibited large concentrated moments at the distal end were impractical to wear due to discomfort. As such, a secondary objective was defined to minimize such uncomfortable loading, based on the contribution on total compensation moment *M*_*G*_ from the concentrated distal moment *M*_*D*_.

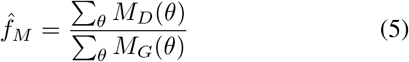

where 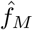represents the normalized contribution of *M*_*D*_ on *M*_*G*_.

#### 2) Particle Swarm Optimization

PSO was chosen for this optimization problem due to its effectiveness in high-dimensional optimization problems [21], and capabilities to be augmented to multi-objective problems [23].

#### a) Single-Objective PSO

For this work, the *particleswarm* MATLAB function from the *Global Optimization* Toolbox was used. During optimization, each particle vector, as well as cost function evaluation and other quantities such as compensation moment and orthosis deformation, were logged to a remote repository such that results could be examined without halting the optimization. Specific details regarding coefficient values used for particle update are given in Appendix D. More information on the use of single-objective PSO for this problem can be found in [16].

#### b) Multi-Objective PSO (MOPSO)

In the context of multi-objective optimization, a solution **x**_**1**_ dominates a so lution **x**_**2**_ if *f*_*i*_(**x**_**1**_) *<*= *f*_*i*_(**x**_**2**_) for all objectives *i*, and there exists at least one objective *j* for which *f*_*j*_(**x**_**1**_) *< f*_*j*_(**x**_**2**_). The set of non-dominated solutions is known as the Pareto set (or Pareto frontier) of the multi-objective optimization problem, and demonstrates the ideal trade-off between the problem’s multiple objectives.

PSO must be modified to consider multiple objectives, as there is no true optimal to define as leaders or personal bests; further, modifications were made to promote diversity among discovered solutions [23]. In this work, we used the *MOPSO* MATLAB function in [24], based on methods in [25] and [26]. Specifically, the population *global best*, a solution typically used to guide all future iterations as it is the historically best solution in single-objective, is instead defined using a weighted probability selection of all Pareto solutions at any given iteration - weighting for this process is defined based on solution isolation from other Pareto solutions. Further, the particles’ *personal bests*, which in single-objective optimization are defined as the historically best solution for each specific particle, are instead defined based on dominance i.e., the logged particle best is the solution that dominates previous particle-specific solutions; if there is no dominance between the currently evaluated particle solution and its previous best, the solution to be retained is selected randomly. Finally, this algorithm implements mutations to mitigate against local minima and promote solution diversity.

#### 3) Non-Linear Constraints

To promote feasible solutions that adhered to specific design constraints, nonlinear constraints were implemented into the optimization process. As nonlinear constraints are not available in PSO, they must be implemented through penalty costs. In our work, each solution that violated a nonlinear design constraint was artificially assigned an objective score of 200 % for all objectives; for comparison, an orthosis design that provides 0 Nm throughout the entire range of motion (ROM) would result in an 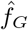 of 100 %. Where possible, nonlinear constraints were evaluated prior to simulation to save computational time. Below is a list of the main design constraints that were considered in our optimization framework. More information on each constraint is given in Appendix D.

1. *Undeformed Intersections*: intersections between any edge defining the orthosis contour in its undeformed state.
2. *Deformed Intersections*: intersections between any edge defining the orthosis contour in its fully deformed state.
3. *Incomplete Rotations*: failures to complete full rotations based on simulation results, typically indicative of device failure; in our work, we retained all solutions that completed rotations of 165° or greater.
4. *Initial Collisions*: overlap between the undeformed orthosis and a region representing the user’s arm at 0°.
5. *Shoulder Collision*: overlap between the fully deformed orthosis and a region representing the user’s shoulder.
6. *Final Collisions*: overlap between the fully deformed orthosis and a region representing the user’s arm the final rotation.
7. *Printer Bounds*: orthosis footprint exceeds bounds corresponding to the print bed of the Roboze INC. Argo 500 (X and Y dimensions of 0.5 x 0.5 m).

### F. Optimization Sequence Overview

An investigation into framework settings (optimization type, kinematic compatibility, output moment specificity) was done to determine the simplest combination of computational settings that could identify effective and wearable orthoses. As such, a series of optimizations was ran with increasingly complex configurations, as seen in Table II.

**TABLE II.**
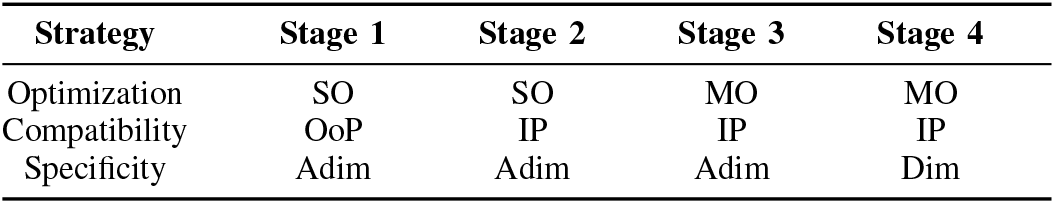
Optimization Configuration by Stage.

Default design parameter bounds are given in Table S1 in Appendix A. Bounds that were altered between optimizations, as well as optimization-specific constraints, are seen in Table III.

**TABLE III.** Optimization Constraints and Bounds by Stage.

| Modification | Stage 1 | Stage 2 | Stage 3 | Stage 4 |
| --- | --- | --- | --- | --- |
| **Constraints** |  |  |  |  |
| Undef. Int. | ✓ | ✓ | ✓ | ✓ |
| Def. Int. | ✓ | ✓ | ✓ | ✓ |
| Incomplete Rot. | ✓ | ✓ | ✓ | ✓ |
| Init. Col. | × | ✓ | ✓ | ✓ |
| Shoulder Col. | × | ✓ | ✓ | ✓ |
| Final Col. | × | ✓ | ✓ | ✓ |
| Print. Bounds | × | × | × | ✓ |
| **Bound Changes** |  |  |  |  |
| min $t_i$ [cm] | 0.05 | 0.05 | 0.5 | 0.5 |
| $w$ [cm] | 1 | 1 | 1 | [1 5] |
| max $Y_f$ [m] | 0.39 | 0.39 | 0.39 | 0.20 |

**TABLE IV.**
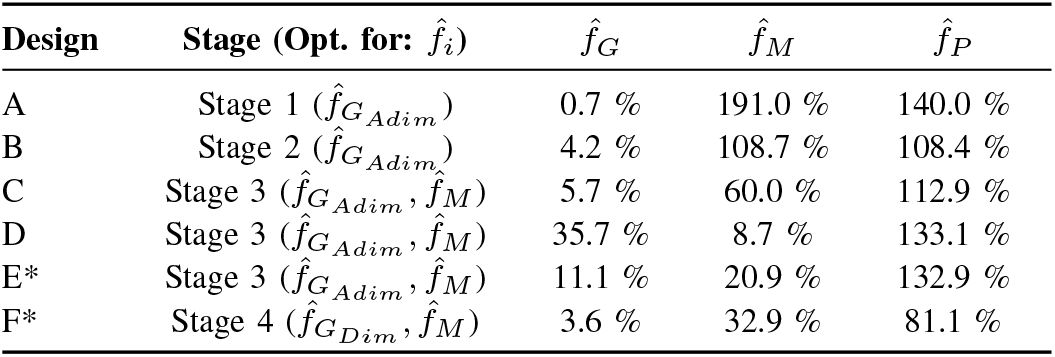
Orthosis Designs by Stage.

#### 1) Stage 1: Single-Objective, Out-of-Plane, adimensional

The first optimization was run with as many simplifications as possible. As such, it was run for a single optimization objective, 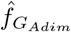, that merely optimized for the shape of the output moment and not the amplitude. Further, no collision constraints between the orthosis and the user were evaluated. This was done to simply understand if, within the encoding definitions provided in our work, it was possible to achieve sinusoidal moment profiles. As this was the first stage of optimization, no initial solutions were inputted, and thus the initial generation of particles were randomly selected from within design vector bounds.

Nonlinear constraints and specific design parameter bounds are seen in Table III. Notably, as this was an adimensional optimization, out-of-plane width *w* was set as a fixed parameter and not included in the optimization process.

#### 2) Stage 2: Single-Objective, In-Plane, adimensional

The second stage of the optimization was run to investigate if orthosis designs can still produce ideal sinusoid profiles when collisions with the user are considered. Thus, the same optimization objective, 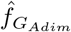, was used as Stage 1, while non-linear constraints were implemented to evaluate for collisions. Nonlinear constraints and specific design parameter bounds are seen in Table III.

Among the initial 50 particles used for this optimization, the 15 best solutions from Stage 1 (corrected for different shapes using the Procrustes analysis) were integrated. Further, an exploratory multi-objective modification of Stage 1 was run prior to Stage 2: 15 solutions selected from the Pareto front of that optimization were also integrated as initial solutions for Stage 2. The remaining 20 particles were generated randomly.

#### 3) Stage 3: Multi-Objective, In-Plane, adimensional

The third stage of optimization was run to investigate how orthosis designs alter between gravity compensation accuracy and a secondary objective. As before, compensation accuracy is optimized by minimizing 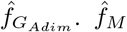 was selected as the secondary objective as large concentrated moments at the distal end of the orthosis limit the operational ROM within which the device can comfortably be worn and used. Nonlinear constraints and specific design parameter bounds are seen in Table III. Notably, the minimum in-plane thickness *t*_*i*_ was increased, as the in-plane thickness at the distal end scaled with distal moment; as such, designs minimizing distal moment yielded extremely small distal thicknesses, which was found to be infeasible upon prototype testing.

Among the initial 50 particles used for this optimization, the 20 best solutions from Stage 2 that were sufficiently different as defined by Procrustes analysis were integrated, while the remaining 30 were randomly generated.

#### 4) Stage 4: Multi-Objective, In-Plane, dimensional

The optimization objectives for this optimizer were 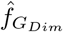, the dimensional objective function targeting a moment profile with amplitude of 8 Nm, and 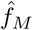. Nonlinear constraints and specific design parameter bounds are seen in Table III. Notably, out-of-plane width *w* was now included as a design parameter; further, the maximum y-coordinate of the distal attachment *Y*_*f*_ was decreased, to only identify solutions that did not attach beyond the user’s elbow, therefore ensuring elbow motion is not constrained and promoting wearability.

Prior to running this multi-objective optimization, multiple iterations of its single-objective equivalent (optimizing for 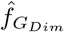) were ran to fine-tune design parameter bounds. While the results of those single-objective optimizations are not shown in this work, the 11 best (differently shaped) solutions from the most recent single-objective optimization were used among the initial 50 particles for this optimization. The remaining 39 particles were randomly generated.

### G. In-Vivo Experiments

#### 1) Orthosis Fabrication

Two orthoses were selected for full-scale in-vivo validation. The primary device, Design E (Fig. 5), was derived from the Stage 3 multi-objective, in-plane, adimensional optimization. It was chosen for its balance between gravity compensation accuracy and low concentrated distal moment. Manufacturing methods, materials, and user interfaces are given in Appendix E.

**Fig. 5.**
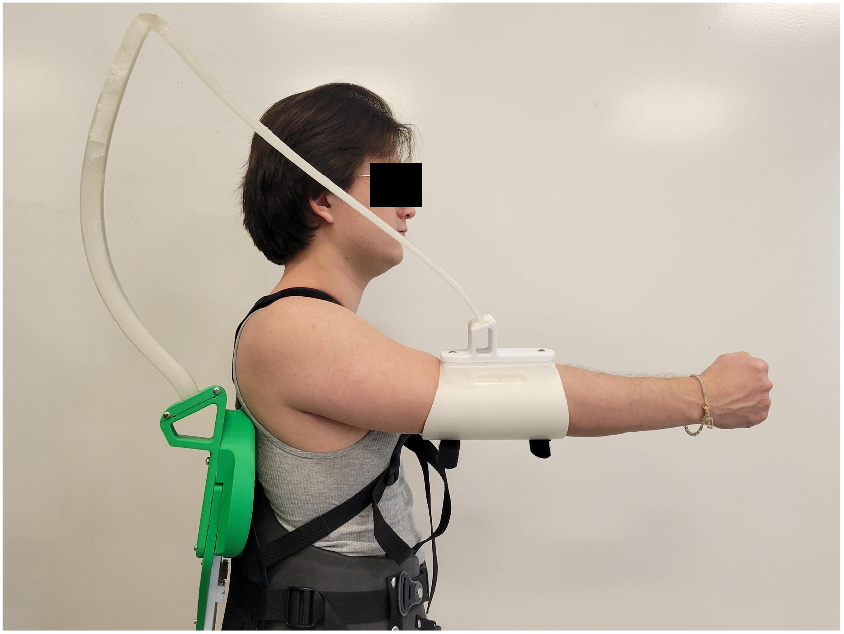
Orthosis based on Design E used for in-vivo testing, worn by one of the authors of this article. The participants don the orthosis at two junctions: 1) a backpack-style harness and 2) an adjustable cuff. 3D printed components tuned to the specific orthosis are used to attach the device to these junctions.

#### 2) Experimental Protocol

The experimental protocol was approved by the University of Delaware Institutional Review Board under IRB #2386926-1. 6 individuals participated in the study (aged 23-34, 5M/1F). Inclusion criteria included no pain or musculoskeletal injuries affecting arm function. The primary experiment solely used the Design E orthosis. A single participant also returned to perform the same experiment with second selected orthosis (Design F, Fig. S11), to determine if shoulder support can still be achieved while leaving the elbow free to move.

The study evaluated the effect of the orthosis on muscles contributing to shoulder flexion during static tasks within ROM where gravitational shoulder moment is large. Additional ROM, muscles, and dynamic tasks were also assessed. Muscles analyzed were anterior deltoid (AD), pectoralis major (PM), upper trapezius (UT), and posterior deltoid (PD). The AD is the primary shoulder flexor, the PM and UT provide supporting functions, and the PD acts as an extensor and antagonist to the orthosis-generated moment.

The experimental protocol followed a 2×2 design, with two experimental factors: 1) Orthosis - participants performed an equal amount of movements with and without the orthosis, and 2) Weight - participants performed an equal amount of movements while holding and not holding a 5 lb. dumbbell. The weight was used to elicit stronger signal in agonist muscle activity to more precisely identify the effect of wearing the orthosis.

The experimental protocol progressed as follows: participants performed four rounds of movements, with each round consisting of 1) a static task, 2) a slow dynamic task, and 3) a fast dynamic task. Experimental factors were alternated to cover all factor combinations in the following order: 1) No Orthosis - Weight, 2) Orthosis - Weight, 3) Orthosis - No Weight, and 4) No Orthosis - No Weight.

For static tasks, participants held a series of static poses, starting with the arm fully raised (*θ* = 0°), and lowered their arm at increments of 30° until reaching 120°, at which point they repeated the task in reverse. Each pose was held for at least 3 seconds, with the 120° being held for twice as long. Participants were asked to keep their elbow extended for all holds.

For dynamic tasks, participants moved between shoulder positions corresponding to shoulder angles of 60° and 120°. Participants began at 60°, moved to 120°, and returned to 60°, moving as naturally as possible, without stopping. Participants repeated this sequence for 5 times total. Participant movements were timed to a metronome, such that they were asked to reach each pose (60°, 120°) on time with a metronome tone. Slow dynamic tasks were performed at 40 beats per minute (bpm), while fast tasks were performed at 60 bpm.

Participants performed all tasks beside a set of targets indicating different arm postures (Fig. S11).

Muscle activity was recorded via electromyography (EMG) using Delsys Inc. (Boston, MA, USA) Trigno Avanti sensors (*F*_*s*_ = 1259 Hz), placed in accordance to SENIAM guidelines. For the PM, the sensor was placed near the collarbone [27]. Accelerometer and gyroscope data were also recorded (*F*_*s*_ = 150 Hz).

Participants were also video recorded during movements. This was primarily to perform quality control over experimental execution, but was also use to define start and end times of dynamic trials. One camera was placed to view the sagittal plane of the participant, while another was used to record the frontal plane. Kinovea (a free 2D motion analysis software under GPL v2 license developed in 2009 [28]) was used to process video recordings and extract shoulder joint angle. To synchronize recordings between EMG sensors and video recordings, the PM sensor was tapped by an experimenter in view of the sagittal-camera, allowing the moment of contact to be used for synchronization.

To normalize muscle activity outcomes, maximum voluntary contractions (MVCs) were performed for each muscle. This was done separately for each muscle with an experimenter providing resistance such that the contraction was isometric. For each muscle, three MVCs were collected, with 10-15 seconds of rest in between. Participants were asked to spend roughly 3 seconds ramping up to their maximum contractions, and to hold each maximum contraction for at least 2 seconds.

Finally, to record subjective outcomes relating to participant perception of wearing the orthosis, a subset of the NASA TLX questionnaire was performed [29] at the end of the experiment. Questions are given in Appendix F.

#### 3) Data Analysis

Data processing was performed primarily in MATLAB, with the exception of joint kinematics for dynamic trials, which also included Kinovea in the process workflow.

All EMG muscle activity data was demeaned, before being band-passed using a Butterworth 4th order zero-phase digital filter with passband frequencies of [20 400] Hz. Signals were then rectified before being low-passed with a Butterworth 4th order zero-phase digital filter, with a 6 Hz cutoff frequency.

All accelerometer and gyroscope data was low-passed using a Butterworth 4th order zero-phase digital filter with a 5 Hz cutoff frequency.

MVC values for normalization were extracted for each muscle by applying a 500 ms moving average [30] to the time series of each contraction dataset, which included three MVC trials. The maximum of the moving average was used as the normalization value.

Static trials were processed using the AD gyroscope signal along the sensor’s Z-axis to identify transitions between static poses. A trial-specific threshold was applied to detect movement; time points at which the signal fell below this threshold and remained below for at least 1 s were flagged as stationary periods. A sampling window was initiated at a predefined offset (default: 1 s) following this flag and ended 1 s later. This procedure yielded 10 sampling windows per static trial (two consecutive windows at 120°). Accelerometer data from the AD sensor’s Y-axis were used to confirm that participants remained stationary during the identified periods. Muscle activity within each window was normalized using muscle-specific MVC values and then averaged.

Dynamic trial processing was applied to both slow and fast trials. Kinematic signals from the Kinovea markerless motion capture were used to define trial windows. Videos were imported into Kinovea and cropped to begin at the PM sensor tap. Virtual markers were placed at the approximate axis of shoulder flexion rotation, the right wrist, and the user’s hip, roughly at the iliac crest. The angle defined by these points was used as shoulder angle. The resulting joint kinematics were outputted as a CSV file for subsequent analysis. Shoulder angular velocity was estimated as the numerical derivative of shoulder angle. A trial-specific threshold (default: *±* 0.3°*/s*) was used to define a deadband on this velocity profile, which enabled the identification of start and stop points for flexion (arm moving up) and extension (arm moving down) trials. Similar to static trials, muscle activity within each window was normalized by muscle-specific MVCs. Averages over each window were determined, and then averaged again over movement type (flexion vs. extension), for each participant.

Linear mixed models for each muscle were used to evaluate the effect of wearing the orthosis on muscle activity. For static trials, a four-way mixed model was performed for each of the muscles with the following factors: 1) Orthosis (2 levels: NoOrtho, Ortho), 2) Weight (2 levels: NoWeight, Weight), 3) Posture (5 levels: 0:30:120), 4) Repetition (2 levels: 1, 2). The mixed model incorporated Participant as the random effect, and included all interaction terms between fixed effects. In the event of a significant effect of any Orthosis-related factor or interaction, post-hoc pairwise comparisons of Orthosis levels were performed using Student’s t-tests with Bonferroni correction for comparisons within each level of Weight, Posture, and Repetition. Effect sizes are reported as the standardized mean difference (Cohen’s *d*_*Z*_).

Towards a more practical assessment of the effects of wearing the orthosis on muscle activity, a secondary three-way linear mixed model was run on muscle-specific outcomes from static trials. Specifically, postures were determined as either “Functional”, i.e., in the middle of the ROM, or “Extreme” i.e., near the boundary of the ROM. All postures performed at 0° or 30° were labeled as as “Extreme”, while those at 60°, 90°, and 120° were labeled as “Functional”. As such, these linear models were run with the following factors: 1) Orthosis (2 levels: NoOrtho, Ortho), 2) Weight (2 levels: NoWeight, Weight), 3) Posture Type (2 levels: Functional, Extreme). The mixed model incorporated Participant as the random effect, and included all interaction terms between fixed effects. In the event of a significant effect of any Orthosis-related factor or interaction, post-hoc pairwise comparisons of Orthosis levels were performed using Student’s t-tests with Bonferroni correction for comparisons within each level of Weight and Posture Type. Effect sizes are reported as the standardized mean difference (Cohen’s *d*_*Z*_).

Finally, four-way linear mixed models were also ran for dynamic trials for each muscle using the following factors: 1) Orthosis (2 levels: NoOrtho, Ortho), 2) Weight (2 levels: NoWeight, Weight), 3) Movement (2 levels: Flexion, Extension), and 4) Speed (2 levels: Slow, Fast). The mixed model incorporated Participant as the random effect, and included all interaction terms between fixed effects. In the event of a significant effect of any Orthosis-related factor or interaction, post-hoc pairwise comparisons of Orthosis levels were performed using Student’s t-tests with Bonferroni correction for comparisons within each level of Weight, Movement, and Speed. Effect sizes are reported as the standardized mean difference (Cohen’s *d*_*Z*_).

For subjective outcomes from the NASA-TLX questionnaire, responses for each domain were averaged across participants.

## III. Results

### A. Optimization

#### 1) Stage 1: Single-Objective, Out-of-Plane, adimensional

This optimization was run for 58 iterations, evaluating 2,900 orthosis designs. The optimizer was stopped as the overall best solution was not improved by greater than 0.5 % (absolute improvement) in 23 iterations (Fig. S8). The global best of this optimization, hereafter referred to as Design A, had a gravity compensation error 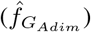 of 0.7 %. This solution heavily relied on concentrated distal moments to achieve its level of gravity compensation (Fig. 6, left). Further, Design A is a very large design that overlaps with the user upon deformation (Fig. 6, second column).

**Fig. 6.**
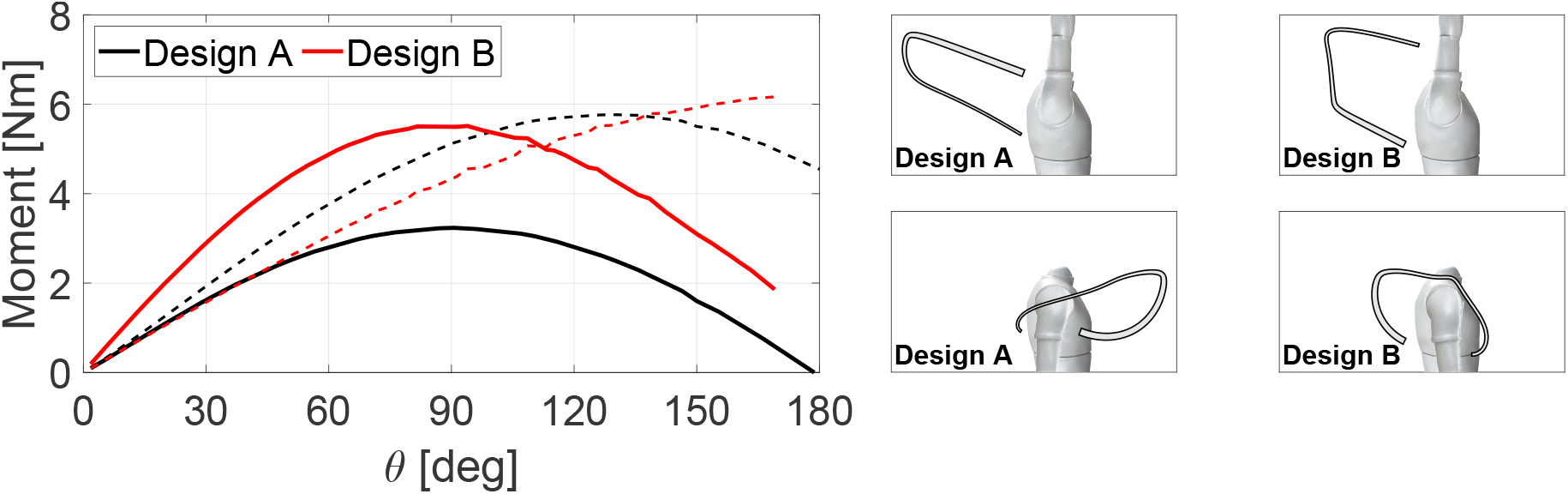
Stage 1 and 2 optimization results. Left: moment profiles for Designs A (black lines - output of Stage 1) and B (red lines - output of Stage 2). Solid lines represent the compensation moment profiles, while dashed lines represent the concentrated distal moments. The middle column shows the undeformed and deformed configurations of Design A, while the right column shows undeformed and deformed configurations of Design B.

#### 2) Stage 2: Single-Objective, In-Plane, adimensional

This optimization was run for 188 iterations evaluating 9,400 orthosis designs. The optimizer was stopped as the overall best solution was not improved by greater than 0.5 % (absolute improvement) in more than 150 iterations (Fig. S8). This optimization was run longer than the previous stage towards identifying a solution with similar compensation accuracy. At best, this optimization outputted a solution (hereafter referred to as Design B) with a compensation error of 4.2 %, a solution more than 5 times worse than the previous stage, identified after running the optimization more than 3 times as long. This design remains large, but is able to wrap around the user (Fig. 6, third column).

#### 3) Stage 3: Multi-Objective, In-Plane, adimensional

This optimization was run for 200 iterations, evaluating 10,000 orthosis designs. The Pareto-extreme solution with best 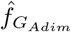, Design C, yielded a gravity compensation score of 5.7 %, similar to the best score of the last stage. This solution exhibited a very large distal moment profile. In contrast, the Pareto-extreme solution with best 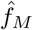, Design D, yielded a distal moment score of 8.7 %; while the distal moment profile remained small, the compensation profile was not at all sinusoidal. The solution selected for in-vivo testing, Design E, yielded 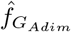 and 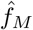 scores of 11.1 % and 20.9 % respectively. These designs can be seen in Fig 7. Multi-objective-based convergence metrics are given in Appendix D.

**Fig. 7.**
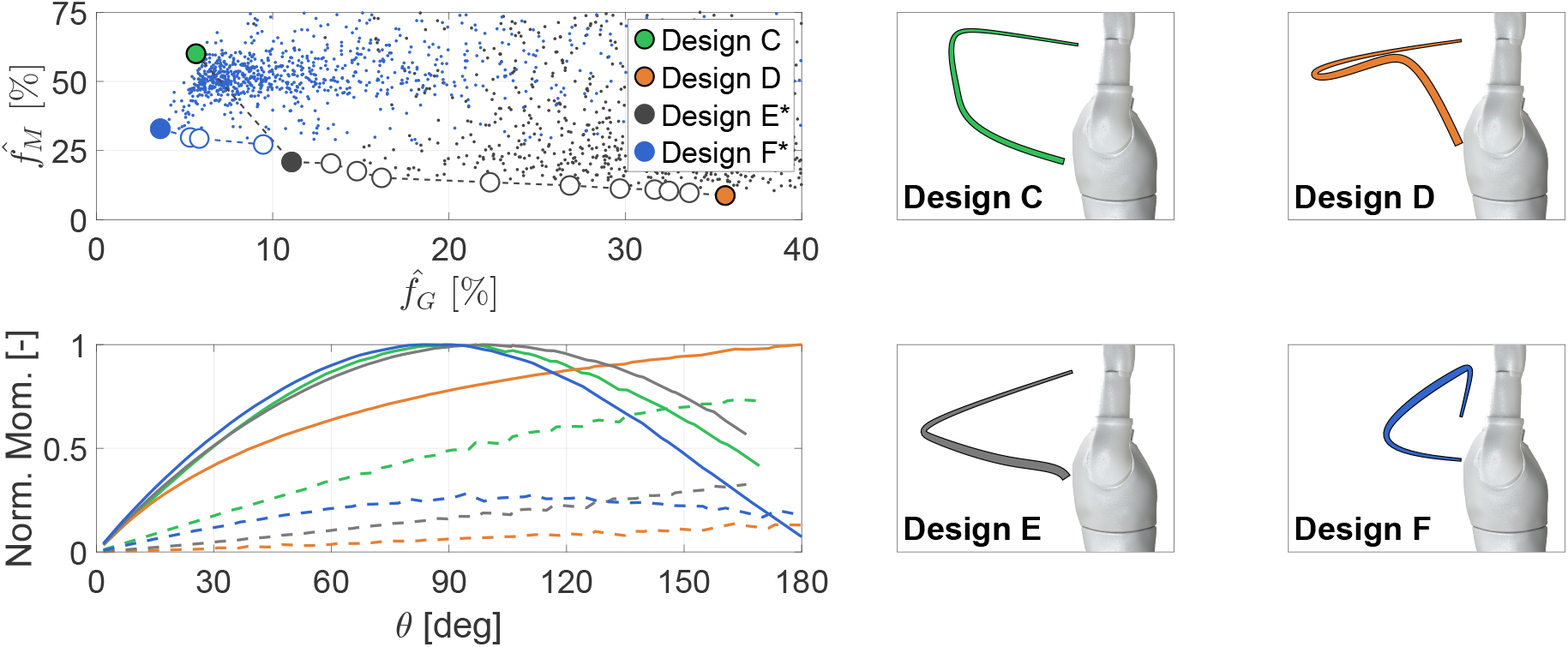
Stage 3 and 4 optimization results. The top left plot visualizes orthosis solutions in optimization objective space, plotted per their optimization objective scores. Results from Stage 3 are plotted in black, while those from Stage 4 are plotted in blue. Pareto solutions are visualized as filled circles. Designs C and D, the two Pareto-extreme solutions from Stage 3, are highlighted in green and orange respectively. Designs E and F, the two designs selected for in-vivo testing from Stages 3 and 4, are colored in black and blue respectively, and are further annotated with asterisks. The bottom left plot shows moment profiles for Designs C, D, E, and F, color-coded per their corresponding highlights. Solid lines represent compensation moment profiles, while dashed lines represent concentrated distal moments. The undeformed configurations of each of these four designs are seen in the middle and right columns.

#### 4) Stage 4: Multi-Objective, In-Plane, dimensional

This optimization was run for 198 iterations, evaluating 9,900 orthosis designs. In this case, 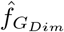 was used for the gravity compensation objective. This optimization heavily favored identifying solutions of low 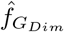 and relatively high 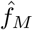; further, all Pareto solutions are similarly shaped. The best orthosis design relative to gravity compensation accuracy, Design F, was selected for in-vivo testing, and yielded 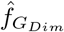 and 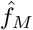 scores of 3.6 % and 32.9 % respectively, producing improved gravity balancing over all solutions outputted in Stage 3 (Fig 7, top left).

### B. In-Vivo Testing

The four-way static mixed model returned a significant effect of Orthosis on AD muscle activity (*F* (1, 195) = 40.07, *p <* 0.0001) (Fig. 8, top row). This was primarily driven by reductions for postures at 60° and 90°. A significant relative reduction of 32 % in muscle activity was present in the Weight condition during the second repetition of the 60° posture (NoOrtho vs Ortho mean ± sem [%MVC]: 58.73 ± 4.51 vs. 40.12 ± 4.51, *p* = 0.0002, *d*_*Z*_ = 0.9639). The four-way mixed models also returned significant effects of Orthosis across the other three muscles (Fig S12, *p <* 0.0001 for all three of PM, UT, and PD).

**Fig. 8.**
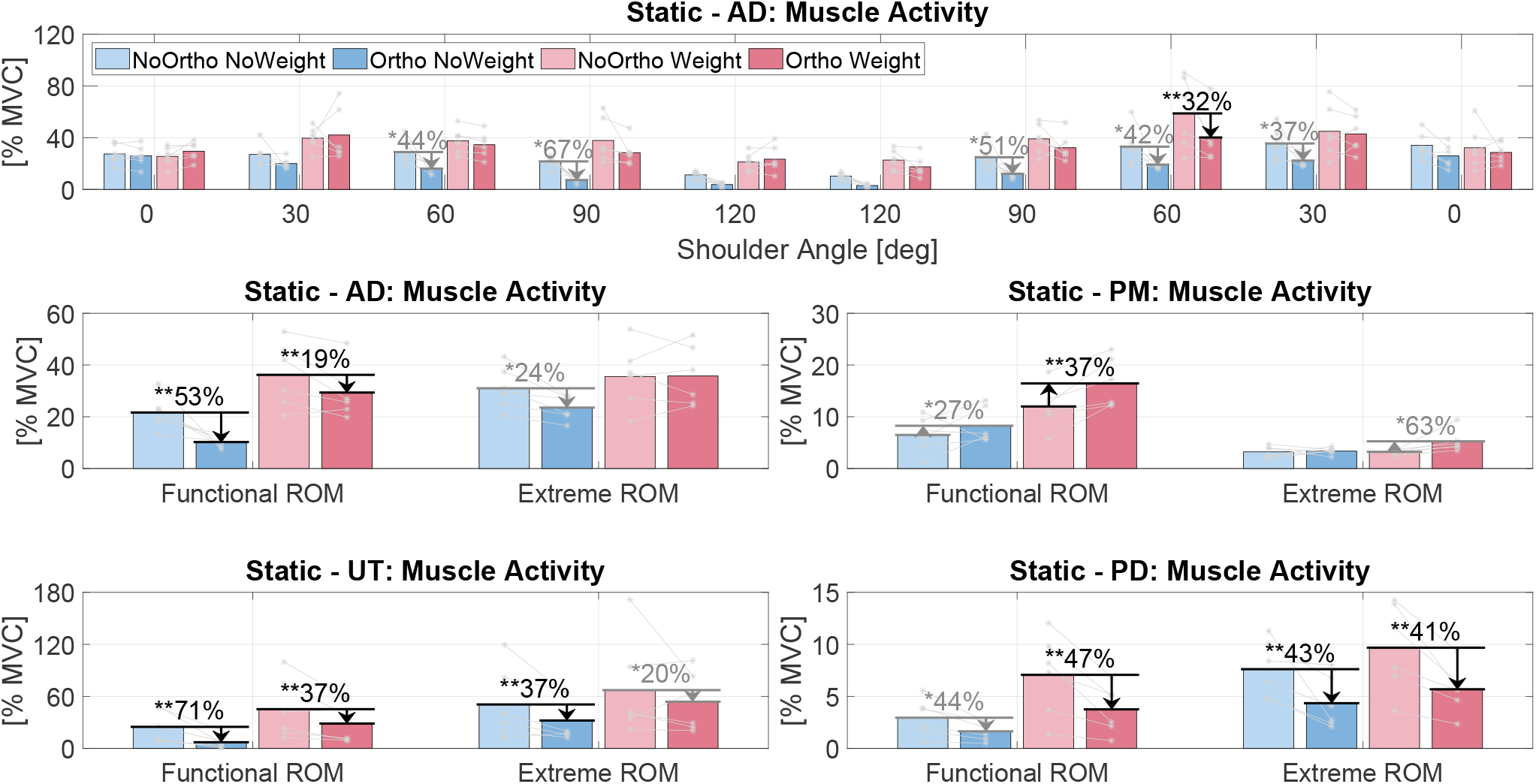
Muscle activity measured during static tasks. Top row: breakdown of AD muscle activity by shoulder angle and posture repetition. Individual participant averages are marked by asterisks, and joined via lines. Arrows and numbers indicate significant relative changes in muscle activity due to wearing the orthosis. Black arrows/text indicate significance after Bonferroni correction, and grey indicates significance uncorrected for multiple comparisons. Bottom two rows: Muscle activity for all muscles, grouped by posture type: Functional vs. Extreme.

The three-way static mixed model for AD returned a significant effect of Orthosis on muscle activity (*F* (1, 227) = 18.79, *p <* 0.0001, Fig. 8, middle row). This was primarily driven by the Functional Posture Type, with significant relative reductions of 53 % and 19 % for the NoWeight (21.64 ± 3.40 vs. 10.21 ± 3.40, *p <* 0.0001, *d*_*Z*_ = 1.4088) and Weight (36.20 ± 3.40 vs. 29.35 ± 3.40, *p* = 0.01, *d*_*Z*_ = 1.0602) conditions respectively.

The three-way static mixed model for PM also returned a significant effect of Orthosis on muscle activity (*F* (1, 227) = 24.30, *p <* 0.0001, Fig. 8, middle row). This was primarily driven by a significant increase of 37 % in the Weight Functional condition (11.98 ±1.02 vs. 16.44 ±1.02, *p <* 0.0001, *d*_*Z*_ = 1.6663).

The three-way static mixed model for UT returned a significant effect of Orthosis on muscle activity (*F* (1, 227) = 36.45, *p <* 0.0001, Fig. 8, bottom row), primarily driven by significant relative reductions of 71 % and 37 % in NoWeight (24.85 ± 11.34 vs. 7.10 ± 11.34, *p* = 0.0004, *d*_*Z*_ = 1.2407) and Weight (45.27 ±11.34 vs. 28.65 ±11.34, *p* = 0.0008, *d*_*Z*_ = 0.9057) Functional conditions respectively, but also a significant relative reduction of 37 % in the NoWeight Extreme condition (50.68 ± 11.60 vs. 32.18 ± 11.60, *p* = 0.0023, *d*_*Z*_ = 0.8806).

Finally, the three-way static mixed model for PD returned a significant effect of Orthosis on muscle activity (*F* (1, 227) = 64.31, *p <* 0.0001, Fig. 8, bottom row). This was primarily driven in the Extreme Posture Type by significant relative reductions by 43 % and 41 % in the NoWeight (7.62 ± 1.07 vs. 4.34 ± 1.07, *p <* 0.0001, *d*_*Z*_ = 1.1783) and Weight (9.68 ± 1.07 vs. 5.68 ± 1.07, *p <* 0.0001, *d*_*Z*_ = 1.5610) conditions respectively, but also by a significant relative reduction by 47 % in the Weight Functional condition (7.07 *±* 1.02 vs. 3.76 *±* 1.02, *p <* 0.0001, *d*_*Z*_ = 1.5535).

The four-way dynamic mixed model for AD returned a significant effect of Orthosis on muscle activity (*F* (1, 75) = 16.56, *p <* 0.0001, Fig. 9, first column). This was primarily driven by NoWeight Flexion movements across Speed types, with relative reductions of 40 % and 34 %, for Slow and Fast trials respectively, that were not significant after correcting for multiple comparisons.

**Fig. 9.**
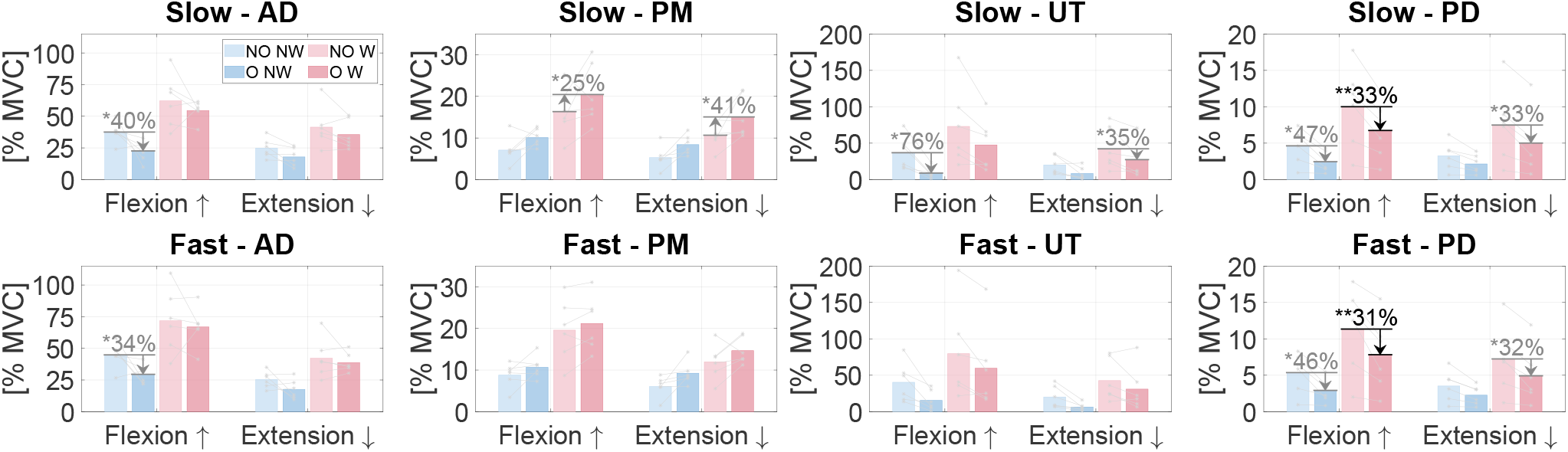
Muscle activity measured during dynamic tasks. Results are plotted by condition combination, movement type, and speed. The columns from left to right show results for AD, PM, UT, and PD in order. Top row shows results for Slow movements, while the bottom row shows results for Fast movements Legend abbreviations: NO - NoOrtho, O - Ortho, NW - NoWeight, W - Weight.

The four-way dynamic mixed model for PM also returned a significant effect of Orthosis on muscle activity (*F* (1, 75) = 23.34, *p <* 0.0001, Fig. 9, second column). This was primarily driven by increases in Weight Slow movements, with relative reductions of 25 % and 41 %, for Flexion and Extension movements respectively, that were not significant after correcting for multiple comparisons.

The four-way dynamic mixed model for UT also returned a significant effect of Orthosis on muscle activity (*F* (1, 75) = 21.24, *p <* 0.0001, Fig. 9, third column). This was primarily driven by Slow movements, with a relative reduction of 76 % observed in NoWeight Flexion movements, and a relative reduction of 35 % observed in Weight Extension movements. Both changes were were not significant after correcting for multiple comparisons.

Finally, the four-way dynamic mixed model for PD also returned a significant effect of Orthosis on muscle activity (*F* (1, 75) = 38.06, *p <* 0.0001, Fig. 9, fourth column). This was primarily driven by significant reductions in Weight Flexion movements, with relative reductions of 33 % and 31 % for Slow (10.01 ±1.51 vs. 6.74 ±1.51, *p* = 0.003, *d*_*Z*_ = 1.7616) and Fast (11.35 ± 1.51 vs. 7.82 ± 1.51, *p* = 0.0014, *d*_*Z*_ = 1.6188) trials respectively.

Tables S2, S3, and S4, given in in Appendix F, show the results of the four-way static, the three-way static, and fourway dynamic linear mixed models respectively. Results to NASA-TLX questions can be found in Table S5.

As mentioned, one participant performed an additional experiment using the Design F orthosis, towards evaluating if reductions in shoulder flexion agonist muscle activity can still be achieved without restraining the elbow. Results are given in Appendix F. Within Functional, NoWeight static trials, relative reductions of 35 % and 59 % were observed in the AD and UT respectively (Fig. S13). Among NoWeight Flexion movements in dynamic trials, no decreases of larger than 10 % were observed in AD; however relative reductions of 36 % and 30 % were observed for Slow and Fast movements in UT muscle activity (Fig. S15).

## IV. Discussion

In this work, we formulated a multi-objective optimization framework for the design of a gravity balancing shoulder orthosis capable of accounting for multiple design objectives spanning both functionality and wearability. Due to the complexity of the real-world problem, a number of simplifications were made to the optimization problem. To investigate the impact of each introduced simplification, we ran a sequence of optimizations, making the optimization problem progressively more complex. Optimization results between sequence stages were compared. Select orthosis designs from this sequence were chosen for in-vivo prototype testing, where the effect of wearing the orthosis was evaluated on muscle activity for select muscles during static and dynamic tasks.

### A. Optimization

When accounting for in-plane collisions as a design constraint, our framework established that it was possible to achieve acceptable levels of shoulder compensation while preventing collisions between orthosis and user. Indeed, as is shown in Fig. 6, the results of Stage 2, which differs from Stage 1 solely by collision constraints, can compensate the shoulder with only 4.2 % error; however, as mentioned, this compensation error is almost an order of magnitude worse than the hypothetical compensation error produced by the output of Stage 1: 0.7 %. While this reduction in accuracy is expected, prior to this work it was unclear to what degree this score would worsen.

While both designs from Stage 1 and 2 are large, the Stage 2 output attached farther along the arm, such that collisions with the user do not occur as the device is deformed. This suggests that attaching orthoses further along the arm may be a strategy towards inhibiting collisions, but perhaps increase compensation error the farther this attachment is. Finally, it is seen that both designs require large distal moments to yield their respective compensation moments.

Upon the implementation of a secondary objective in the optimization, orthosis designs remain similar to the Stage 2 output in cases where compensation accuracy is still prioritized (Designs B vs. C). Progressing along the Pareto front of Stage 3, it is seen that the optimal orthosis design shape changes as compensation accuracy is prioritized less, in favor of minimized distal moment (Fig. 7). Specifically, the gentle, two-corner design seen in Design C is altered in favor for a single, harsher bend in Design E. As priority shifts more to 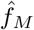, the angle of this single bend continues to decrease, until it is almost non-existent in Design D. The Stage 3 optimization seemed to favor investigating regions of low distal moment, as evidenced by the clustering of solutions (black) seen in Fig. 7. In fact, the Pareto design with the best 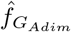, Design C, was extremely isolated. As such, for in-vivo testing, the design with the next best 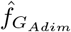 (Design E) was selected due to the large improvement in 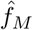.

Stage 4 extended Stage 3 by pursuing a dimensional optimization that controlled for printability and exact arm attachment locations. In Stage 4, the optimization thoroughly investigated a constrained region of the solution space, producing a Pareto front with low 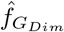 (3.6 %), but did not make large improvements in terms of 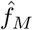. In fact, all Pareto solutions were slightly different encodings corresponding to the same design; as such, the Pareto solution with the best 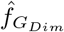was selected for in-vivo testing (Design F).

Thus, opposite regions of the true Pareto front were explored in the runs for Stages 3 and 4, with Stage 3 exploring solutions with improved distal moments, while Stage 4 discovered improved gravity compensation. This clustering of discovered solutions suggests only partial Pareto exploration from either optimizer. While the MOPSO algorithm implements multiple features to promote exploration and diversification, namely mutations and the implementation of dominance into particle update, it is still possible that thorough exploration is not achieved. Indeed, in an optimization with many nonlinear constraints, mutating orthosis designs to random solutions inhibits a large subset of particles from identifying useful solutions each iteration. Further, this implementation of MOPSO selects the global best particle in a probabilistic approach, weighting the probability of each Pareto solution to be selected based on its isolation. While strategically sound in attempting to promote a dense Pareto front, it is possible that the evolution of particles within this high-dimensional problem is slow enough to require the global best to be selected deterministically, allowing many iterations with the same leader, thus more effectively drawing solutions towards unexplored regions of the solution space.

As an extension of this work, an exploration into the scaling of the multi-objective optimization using a size objective (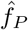 - more information given in Appendix D) as the third objective was performed. Specifically, a direct extension of Stage 4 was performed with three objectives. Results are given in Appendix D. This optimization was run for a similar number of iterations as the multi-objective optimizations presented here; while identified solutions performed similarly in 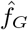, scores in 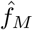 were at best around two times worse than those of Stage 4 (Figs S9 and S10). It is possible that as more objectives are incorporated into this optimization paradigm, the exploration-exploitation tradeoff of the optimizer becomes increasingly complex, thus requiring more iterations to be run for the Pareto front to be exhaustively searched.

### B. In-Vivo Testing

The main goal of the in-vivo testing was to establish the effect of wearing the orthosis in static, unweighted tasks on muscles used to support shoulder flexion, specifically in postures where shoulder moment is expected to be largest. Secondary goals involved an investigation on how orthosis affects muscle activity in other nearby muscles, across different postures, and during dynamic tasks.

Starting with the effects on the shoulder flexion agonists (AD and UT), our study indicates that muscle activity was significantly reduced when participants were wearing the orthosis, by 53 % and 71 % for AD and UT respectively, in the targeted conditions. This is in line with expectations: within the Functional ROM, shoulder moment is at its largest, and hence, where the orthosis is designed to provide the most compensation. At Extreme ROM, a non-significant decrease is seen in the unweighted condition. This is also inline with expectation, as these regions require less support due to small shoulder angles. Further, changes in EMG are difficult to detect at this range, due to noise in measurements associated with the smaller activation required to balance the arm’s gravity load in these configurations. In Weighted conditions, within-participant reductions were smaller, and across-participant variance was larger. While the implementation of Weight trials were done to increase signal in select muscle activity, it is possible that this resulted in increased activity in all muscles due to cocontraction. As such, a significant decrease in AD muscle activity is seen in Functional ROM, again due to this ROM being the main target of support, while no effect is present in Extreme ROM.

The posterior deltoid PD was also included in this experiment as a primary antagonist to shoulder flexion. Changes in muscle activity in the antagonist could provide an indication if overcompensation was being provided by the orthosis. Surprisingly, in targeted conditions, a non-significant decrease of 44 % with orthosis was observed in PD. This effect could possibly indicate a decreased need for joint stabilization for static tasks: in other words, as the orthosis is offloading the primary agonist AD, there is also less of a need for PD to be used to maintain static postures. This stabilization rationale is supported in other factor levels, as significant decreases in muscle activity are observed in Extreme ROM and in Weight conditions, where greater stabilization is maybe necessary.

The pectoralis major PM was also included in this experiment, due to its close proximity and relationship with shoulder motion. Among its functions, the PM is involved in horizontal flexion, which moves the shoulder and arm within the transverse plane. Within targeted conditions, PM muscle activity increased by 27 %, although this increase was not significant after correction, with greater or more significant increases being observed across Weight conditions. It is believed that this increase is associated with lateral motion being caused by the orthosis: as the orthosis was designed and simulated in a 2D plane, out-of-plane motion and forces were not accounted for. Indeed, it was observed during prototype testing that in cases where the device was made from a softer material, or simply too thin, large lateral deflections occur as the orthosis was deformed. While this was largely resolved for in-vivo testing by ensuring large enough out-of-plane width, some lateral deflection still occurred. As such, it is believed that increases in the PM associated with the orthosis were due to participants using this muscle to prevent lateral motion, with this effect being larger when holding a weight, perhaps as the weight may increase instability in the lateral direction.

Among dynamic trials, effects follow similar trends as static cases, but generally to smaller magnitudes. This is somewhat expected as muscular dynamics differ between static and dynamic postures - namely, the sinusoid nature of the arm moment due to gravity is defined based on a series of static postures over the shoulders ROM. Decreases in shoulder flexion agonists were hypothesized to occur during flexion movements in unweighted conditions. Indeed, non-significant decreases were observed in both Slow and Fast movements for AD, and Slow movements for UT. Decreases are observed in other conditions for these agonists, although these decreases are either not-significant after correction, or not significant at all. Further, similar to static trials, decreased muscle activity in PD was observed across conditions. Notably, decreases of more than 30 % were observed in weighted extension movements. While the PD is the antagonist of shoulder flexion, it is the primary agonist for shoulder extension, although the muscle is supported by gravity in lowering the arm. Reductions across experimental conditions indicate to the prior theory: that this muscle is used for stabilization, which is less needed while wearing the orthosis.

Finally, similar to static cases, increases in PM are observed in some weight conditions. As speculated above, it is believed that this is due to lateral motion caused by the orthosis, increased when the user is holding the weight.

The in-vivo validation using Orthosis F, a product of the Stage 4 optimization, demonstrated smaller reductions in shoulder flexion agonists. Upon experimentation, it was clear that this prototype was much softer than Orthosis E, despite being designed to compensate the same moment amplitude. As such, it is expected that effects are less in Orthosis F than E; specifically, this difference is expected to be due to differences in manufacturing processes, rather than due to different constraints at the elbow.

## V. Conclusions

In this work we developed a multi-objective optimization framework for the design of soft, gravity balancing shoulder orthoses that accounts for both functional performance and wearability constraints. We used this framework across multiple rounds of orthosis design optimization, and selected multiple outputs to be prototyped using thermoplastic polyurethane. Prototypes were evaluated on muscle activity surrounding the shoulder, where associated reductions in the anterior deltoid and upper trapezius were observed, as well as in the posterior deltoid. Significant increases in muscle activity were observed in the pectoralis major. These findings demonstrate the framework’s ability to accurately navigate wearability constraints while still identifying optimal orthosis designs, though improvements in simulating out-of-plane loads and refinement into 3+ optimization objectives are needed to fully translate the presented framework into a practical tool to assist the design of gravity balancing shoulder orthoses.

## VI. Acknowledgments

This work is supported by the National Science Foundation under Award 194712. We would like to thank Dr. Suresh G. Advani at the Department of Mechanical Engineering and Center for Composite Materials at the University of Delaware, for providing access to the Argo 500 3D printer, without which fabrication of in-vivo scaled prototypes would not have been possible. We would also like to acknowledge Roboze INC. for their assistance in using the printer.

## References

[1] M. Tschiersky, E. E. G. Hekman, D. M. Brouwer, and J. L. Herder, “Gravity Balancing Flexure Springs for an Assistive Elbow Orthosis,” IEEE Transactions on Medical Robotics and Bionics, vol. 1, no. 3, pp. 177–188, 2019.

[2] M. Tschiersky, E. E. Hekman, J. L. Herder, and D. M. Brouwer, “Gravity Balancing Flexure Spring Mechanisms for Shoulder Support in Assistive Orthoses,” IEEE Transactions on Medical Robotics and Bionics, vol. 4, no. 2, pp. 448–459, 2022.

[3] Z. Cheng, S. Foong, D. Sun, and U. X. Tan, “Towards a multi-DOF passive balancing mechanism for upper limbs,” IEEE International Conference on Rehabilitation Robotics, vol. 2015-Septe, pp. 508–513, 2015.

[4] M. Asgari, E. A. Phillips, B. M. Dalton, J. L. Rudl, and D. L. Crouch, “Design and Preliminary Evaluation of a Wearable Passive Cam-Based Shoulder Exoskeleton,” Journal of Biomechanical Engineering, vol. 144, no. 11, nov 2022.

[5] T. M. Sukal and J. P. A. Dewald, “Following Hemiparetic Stroke : Neuroscientific Implications,” vol. 183, no. 2, pp. 215–223, 2010.

[6] J. P. Dewald and R. F. Beer, “Abnormal joint torque patterns in the paretic upper limb of subjects with hemiparesis,” Muscle and Nerve, vol. 24, no. 2, pp. 273–283, 2001.

[7] M. D. Ellis, T. Sukal-Moulton, and J. P. Dewald, “Progressive shoulder abduction loading is a crucial element of arm rehabilitation in chronic stroke,” Neurorehabilitation and Neural Repair, vol. 23, no. 8, pp. 862– 869, 2009.

[8] M. D. Ellis, T. M. Sukal-Moulton, and J. P. Dewald, “Impairment-based 3-D robotic intervention improves upper extremity work area in chronic stroke: Targeting abnormal joint torque coupling with progressive shoulder abduction loading,” IEEE Transactions on Robotics, vol. 25, no. 3, pp. 549–555, 2009.

[9] L. Van Engelhoven and H. Kazerooni, “Design and Intended Use of a Passive Actuation Strategy for a Shoulder Supporting Exoskeleton,” 2019 Wearable Robotics Association Conference, WearRAcon 2019, pp. 7–12, 2019.

[10] J. Tian, H. Zhu, C. Lu, C. Yang, Y. Liu, B. Wei, and C. Yi, “A Novel Passive Occupational Shoulder Exoskeleton With Adjustable Peak Assistive Torque Angle for Overhead Tasks,” IEEE Transactions on Biomedical Engineering, vol. 72, no. 2, pp. 734–746, 2025.

[11] J. Kim, J. Moon, J. Yoon, S. Kim, S. Lee, and G. Lee, “Quasi-passive shoulder exoskeleton with enhanced assistance variability to adapt to frequent load changes,” International Journal of Robotics Research, vol. 44, no. 12, pp. 2020–2043, 2025.

[12] T. Rahman, W. Sample, R. Seliktar, M. Alexander, and M. Scavina, “A body-powered functional upper limb orthosis,” Journal of Rehabilitation Research and Development, vol. 37, no. 6, pp. 675–680, 2000.

[13] T. Rahman, R. Ramanathan, R. Seliktar, and W. Harwin, “A simple technique to passively gravity-balance articulated mechanisms,” Journal of Mechanical Design, Transactions of the ASME, vol. 117, no. 4, pp. 655–657, 1995.

[14] R. Sanchez, D. Reinkensmeyer, P. Shah, J. Liu, S. Rao, R. Smith, S. Cramer, T. Rahman, and J. Bobrow, “Monitoring functional arm movement for home-based therapy after stroke,” Annual International Conference of the IEEE Engineering in Medicine and Biology - Proceedings, vol. 26 VII, pp. 4787–4790, 2004.

[15] D. Gijbels, I. Lamers, L. Kerkhofs, G. Alders, E. Knippenberg, and P. Feys, “The Armeo Spring as training tool to improve upper limb functionality in hemiplegic Cerebral Palsy: A pilot study,” 2016 IEEE 2nd International Forum on Research and Technologies for Society and Industry Leveraging a Better Tomorrow, RTSI 2016, no. 2008, pp. 1–8, 2016.

[16] H. A. Chishty and F. Sergi, “A Multi-Objective Simulation-Optimization Framework for the Design of a Compliant Gravity Balancing Orthosis,” Proceedings of the IEEE RAS and EMBS International Conference on Biomedical Robotics and Biomechatronics, no. September, pp. 1440– 1445, 2024.

[17] C. V. Jutte and S. Kota, “Design of nonlinear springs for prescribed load-displacement functions,” Journal of Mechanical Design, vol. 130, no. 8, pp. 0 814 031–08 140 310, 2008.

[18] G. Radaelli and J. L. Herder, “Isogeometric shape optimization for compliant mechanisms with prescribed load paths,” Proceedings of the ASME Design Engineering Technical Conference, vol. 5A, no. July, 2014.

[19] G. Radaelli and J. L. Herder, “A monolithic compliant large-range gravity balancer,” Mechanism and Machine Theory, vol. 102, pp. 55–67, 2016.

[20] B. L. Rijff, J. L. Herder, and G. Radaelli, “An energy approach to the design of single degree of freedom gravity balancers with compliant joints,” Proceedings of the ASME Design Engineering Technical Conference, vol. 6, no. PARTS A AND B, pp. 137–148, 2011.

[21] G. Lindfield and J. Penny, “Particle Swarm Optimization Algorithms,” Introduction to Nature-Inspired Optimization, pp. 49–68, jan 2017.

[22] S. Pheasant and C. M. Haslegrave, “Anthropometric Data,” Bodyspace, pp. 239–279, 2019.

[23] C. A. Coello Coello and M. Reyes-Sierra, “Multi-Objective Particle Swarm Optimizers: A Survey of the State-of-the-Art,” International Journal of Computational Intelligence Research, vol. 2, no. 3, pp. 287– 308, 2006.

[24] V. Martínez-Cagigal, “Multi-Objective Particle Swarm Optimization (MOPSO),” 2026. [Online]. Available: https://www.mathworks.com/matlabcentral/fileexchange/62074-multi-objective-particle-swarm-optimization-mopso

[25] C. A. C. Coello, G. T. Pulido, and M. S. Lechuga, “Handling Multiple Objectives With Particle Swarm Optimization,” IEEE TRANSACTIONS ON EVOLUTIONARY COMPUTATION, vol. 8, no. 3, pp. 256–279, 2004.

[26] M. R. Sierra and C. A. C. Coello, “Improving PSO-Based Multi-objective Optimization Using Crowding, Mutation and η-Dominance,” in International Conference on Evolutionary Multi-Criterion Optimization, vol. 1, 2005, pp. 505–519.

[27] H. Król, G. Sobota, and A. Nawrat, “Effect of electrode position on EMG recording in pectoralis major,” Journal of Human Kinetics, vol. 17, pp. 105–112, 2007.

[28] A. Puig-Diví, C. Escalona-Marfil, J.M. Padullés-Riu, A. Busquets, X. Padullés-Chando, and D. Marcos-Ruiz, “Validity and reliability of the Kinovea program in obtaining angles and distances using coordinates in 4 perspectives,” PLoS ONE, vol. 14, no. 6, p. e0216448, jun 2019. [Online]. Available: https://pmc.ncbi.nlm.nih.gov/articles/PMC6550386/

[29] S. G. Hart and L. E. Staveland, “Development of NASA-TLX (Task Load Index): Results of Empirical and Theoretical Research,” Advances in Psychology, vol. 52, no. C, pp. 139–183, jan 1988. [Online]. Available: https://www.sciencedirect.com/science/chapter/bookseries/abs/pii/S0166411508623869

[30] P. Konrad, The ABC of EMG A Practical Introduction to Kinesiological Electromyography, 2005. [Online]. Available: https://www.noraxon.com

