## Appendices for "Multi-objective optimization-based design of a compliant gravity balancing orthosis: development and validation"

### Appendix A
### Orthosis Definition

#### A. Shape Encoding

The following sections will detail different components of the optimization design vector, which defines the overall shape, thickness, and attachment points of an orthosis. The general form of the design vector is seen in (2). The default design parameter bounds are seen in Table S1.

TABLE S1
Default Design Vector Bounds

| Design Var. | Units | Bounds |
|---|---|---|
| $X_0$ | [m] | [-0.20 -0.15] |
| $Y_0$ | [m] | [-0.40 0] |
| $X_f$ | [m] | [-0.15 -0.11] |
| $Y_f$ | [m] | [0.15 0.39] |
| $l_i$ | [m] | [0.01 0.45] |
| $\psi_1$ | [rad] | $[\frac{\pi}{4}\ \frac{\pi}{2}]$ |
| $\psi_i$ | [rad] | [-2 2] |
| $t_i$ | [m] | [0.0005 0.02] |
| $w$ (constant) | [m] | [0.01] |

*1) Orthosis Attachments:* Parameters $X_0$ and $Y_0$ are used to define the horizontal and vertical coordinates for the proximal end of the orthosis respectively (Fig. S2). Relative to the torso, the horizontal direction is defined as the axis perpendicular to the midline of the wearer, while the vertical direction is parallel to the midline. The bounds on these parameters represent a rectangular region within which the proximal attachment can reside. This region is defined such that the proximal attachment will not overlay with the wearer (i.e., an estimate of the thickness of the user's back is considered when defining these bounds). Further, as attachment components used to attach to the orthosis to the user have their own bulk, extra consideration is necessary when defining the boundaries. Finally, the region is constrained such that the proximal end of the orthosis will never be above the shoulder (Fig. S1).

Similarly, $X_f$ and $Y_f$ are used to define the horizontal and vertical coordinates of the distal end of the orthosis (Fig. S2). In this case, the horizontal direction is defined as the axis perpendicular to the neutral axis of the user's arm, while the vertical axis lays parallel. Similarly to the proximal parameters, the bounds on these parameters represent the rectangular region within which the distal attachment can reside (Fig. S1). The rectangle is defined such that the attachment will not overlay with the arm, and considers the bulk of the attachment components. The distal attachment is constrained to always attach past the shoulder (i.e., on the arm), and may or may not attach past the elbow, depending on what stage of the optimization sequence the process is in.

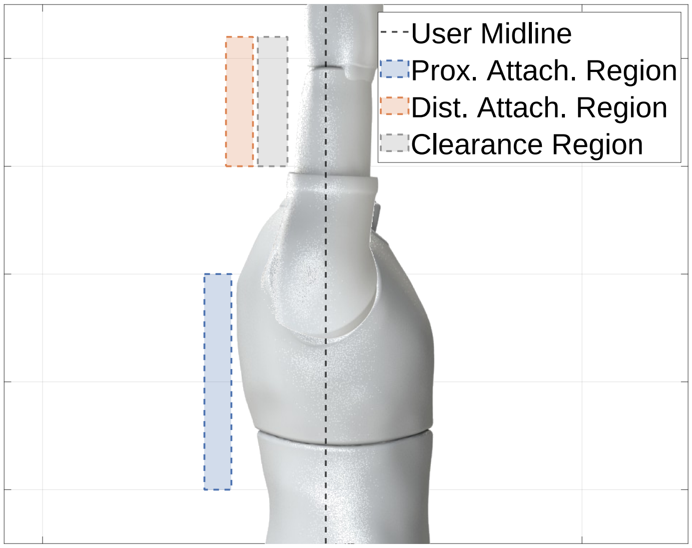


Fig. S1. Orthosis attachment regions. Possible attachment points for the proximal and distal ends of the orthosis are displayed in blue and orange respectively. An example region, gray, is used to portray how the bulk of attachment components needs to be considered when defining attachment regions.

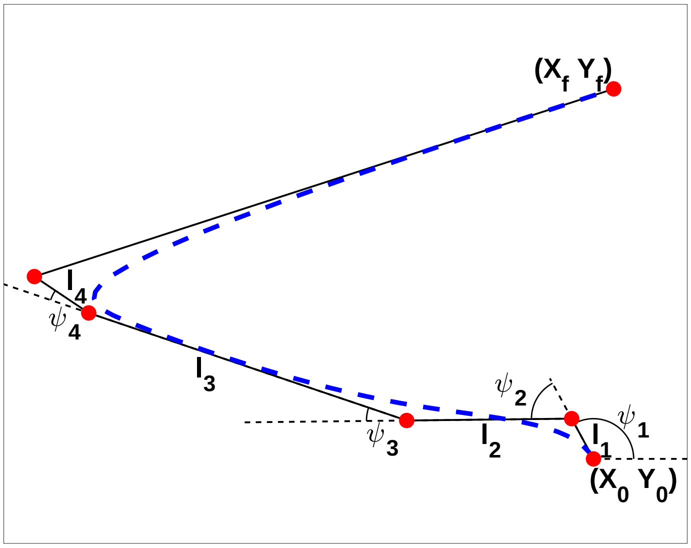


Fig. S2. Neutral axis definition. The neutral axis is a piecewise cubic b-spline defined off a control polygon developed using linkage formulation. $X_0$ and $Y_0$ correspond to the proximal attachment while $X_f$ and $Y_f$ correspond to the distal attachment.

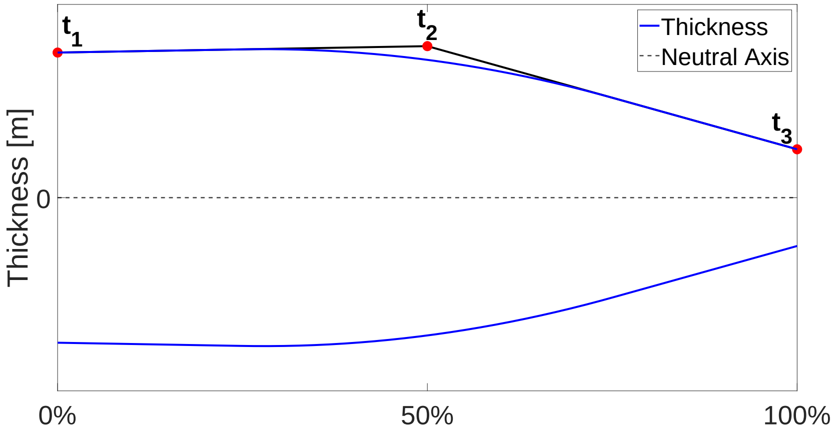


Fig. S3. In-plane thickness profile to define orthosis edges. The profile is defined off of a number of control points, to develop a thickness profile as a function of neutral axis location, using as few parameters as possible. 0 % corresponds to the proximal attachment of the neutral axis, while 100 % corresponds to the distal attachment.

*2) Orthosis Neutral Axis:* The neutral axis of the orthosis is defined using linkage chain formulation. This process begins with a definition of the control polygon, done using the attachment points $X_0$, $Y_0$, $X_f$, and $Y_f$ in conjunction with design parameters $l_1$ to $l_n$ and $\psi_1$ to $\psi_n$. $l_i$ represents the absolute distance from point $i$ of the control polygon to point $i$+1, while $\psi_i$ represents the relative angle at which that $i$th link departs from point $i$. The proximal attachment defined by $X_0$ and $Y_0$ represents the first control point, while the distal attachment defined by $X_f$ and $Y_f$ represents the final point, and is appended to the list of control points defined by the linkage chain formulation (Fig. S2).

The neutral axis is defined off this control polygon as a piecewise cubic b-spline trajectory (the *bsplinepolytraj* MATLAB function). While b-splines do not interpolate all control points, this methodology offers a number of advantages including C2 continuity (for smooth curves), local control (i.e., modifying a specific control point only alters the curve near that point), and an intuitive spatial bound on the curve's shape (i.e., the curve defined by the piecewise spline will entirely reside within the convex hull defined by the control points). This last advantage is a property of bezier curves as a whole.

As b-spline are typically used to define trajectories, when using this methodology to define 2D spatial curves, a trajectory is defined within each dimension relative to time, and then collapsed together using time points to synchronize trajectories.

*3) Orthosis In-Plane Thickness:* Allowing for varying in-plane thickness enables extra customization for orthosis output compensation. As such, $m$ design parameters ($t_1$ to $t_m$) were used to define the in-plane orthosis thickness profile.

The purpose of these parameters was to define the in-plane thickness at each point of the neutral axis (hereby referred to as 0-100 %) with as few design parameters as possible. As such, these parameters were used to define another b-spline (using uniform subdivision of order 3) using the *spcrv* MATLAB function, which provides points along the curve defined in the function input - leveraging another advantage of the bezier curve, that the equation of the curve can be expressed using

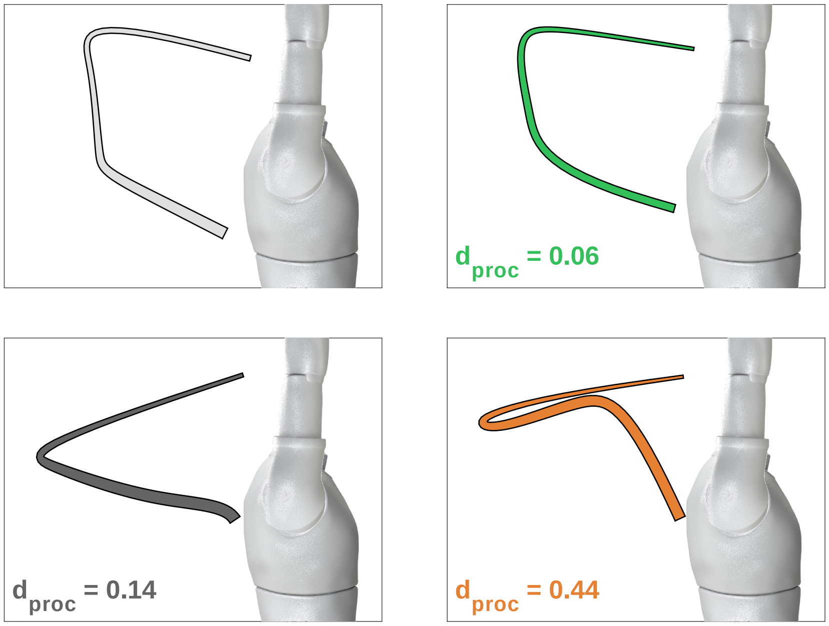


Fig. S4. Procrustes analyses on three orthosis designs (green, black, orange) relative to an initial design (gray). $d_{proc}$ represents the output of the Procrustes analysis: the Procrustes distance. A larger value signifies that the two shapes are more different.

control points. Once the spline was defined, it was interpolated to determine thickness values for each point along the neutral axis (0-100 %). (Fig. S3).

Specifically, this profile represents the half-thickness of the orthosis. Along each point defining the orthosis neutral axis, the perpendicular direction is defined relative to the line connecting the current point to the previous, with the exception of the first point, in which case this definition was done relative to the line connecting the first and second points. Along the perpendicular line, points defining the top and bottom edges are defined at a distance away from the orthosis point defined by the thickness profile. The two sets of points defined by this method define the top and bottom edges of the orthosis respectively, ultimately representing the orthosis 2D contour.

Due to the curvature of the neutral axis, it is possible that the edges of the orthosis self-intersect. In such a case, for each intersection that occurs for a surface, the indices of the line defining the surface are flagged for points that lay within the intersection. These flagged points are removed, and the entire edge is smoothed using a 5-point moving average. This process is repeated until no intersections remain.

*4) Orthosis Out-of-Plane Width:* The final design parameter $w$ defines the out-of-plane width of the orthosis, and is relevant for simulation, as the output loading of the orthosis scales linearly with this parameter. During adimensional optimizations, where scaling is not necessary as a specific moment amplitude is not required, this parameter is omitted as a design parameter, and is instead used as a constant of 1 cm.

#### *B. Shape Similarity*

When examining favorable orthosis solutions for framework output, it is possible that many favorable solutions have similar designs. To increase practicality of the framework, a shape similarity analysis is performed in order to omit solutions that are already present in the output set. This is particularly relevant for the outputs of the framework's single-objective optimizations, where the output set is a collection of differently shaped orthoses with effective gravity compensation, but may also be used in the multi-objective setting.

Using a single-objective case, once the optimization has ended, all solutions (orthosis designs) with compensation errors below a specific threshold (e.g., $<$20%, as in, orthosis designs that under or overcompensates by less than 20%, relative to the selected cost function) are considered for framework output. The solutions are sorted in order of increasing compensation errors (i.e., best to worst). Recursively, all orthosis designs in the set are compared to the best, and those considered too similarly designed are omitted. This process then continues with the second best orthosis design still remaining in the set, and so on.

To evaluate shape similarity between two designs, a Procrustes analysis is performed. This is done by first performing a Procrustes transformation, which is the best shape-preserving Euclidean transformation (consisting of rotation, reflection, scaling, and translation) between the two shapes. This is an optimal transformation that minimizes the sum of squared differences between the landmark points of one shape to the transformed points of the other. The output of the Procrustes analysis of two shapes is the squared error based on the best transformation.

Within the framework, a Procrustes distance of 0.05 or greater was used to denote sufficiently different orthosis designs.

### Appendix B
### Simulations

#### A. Moment Calculations

The results of the nonlinear simulation return nodal forces (as well as displacements) at each load step of the deformation. As such, a compensation moment profile as a function of shoulder angle can be determined. These results are eventually used to calculate optimization objective functions.

Fig. S5 shows three free-body-schematics detailing interactions between the orthosis, arm, and the torso. Beginning with the orthosis and the arm Fig. S5A, the output loading at the distal end of the orthosis relates to the moment felt at the shoulder from the orthosis (i.e., the compensation moment) as:

$$M_{S_A}(\theta) = M_{D_{O-U}}(\theta) + r_{D/S} \times F_{D_{O-U}}(\theta) \tag{6}$$

that is, at angle $\theta$, the moment at the shoulder based on the arm's dynamics $M_{S_A}$ can be determined from the concentrated moment that the orthosis imposes on the user at the distal location $M_{D_{O-U}}$ and the output force the orthosis imposes on the user at the distal location, $F_{D_{O-U}}$.

In practice, nodal forces were directly used to calculate the moment at the shoulder, rather than first being used to calculate concentrated moments and output force; this is done by summing the moment contribution of each nodal force on the distal edge of the orthosis relative to the shoulder:

$$M_S(\theta) = \sum_i r_{i/S} \times F_i(\theta) \tag{7}$$

where $i$ indexes over all the nodes of interest, $r_{i/S}$ represents the distance of node $i$ to the shoulder $S$, and $F_i(\theta)$ is the nodal force of node $i$ at angle $\theta$.

While the above calculation is used to calculate shoulder moment, the discrepency between concentrated moments and output forces is relevant for other optimization objectives, specifically when minimizing the concentrated moment at the distal end.

Unfortunately, simulated nodal force results at the distal end of the orthosis were highly volatile and assumed to be incorrect, possibly due to the rotating frame of the coordinate system fixed to the orthosis edge over time. As such, nodal forces from the proximal end of the orthosis, which did not rotate as it was fixed, were used instead, to determine the moment at the shoulder based on the torso's dynamics $M_{S_T}$, which is the negative of $M_{S_A}$. The rest of this subsection serves to provide the proof that the two shoulder moments are indeed opposites, and therefore using the proximal end of the orthosis to solve for overall compensation is in fact valid.

Fig. S5B shows a free-body-schematic of the orthosis, relating the output loads of each end of the orthosis.

$$F_{D_{U-O}}(\theta) + F_{P_{U-O}}(\theta) = 0 \tag{8}$$

$$M_{P_{U-O}}(\theta) + M_{D_{U-O}}(\theta) + r_{D/P} \times F_{D_{U-O}}(\theta) = 0 \tag{9}$$

Substituting (6) into (9) ultimately yields:

$$M_{S_A}(\theta) + M_{P_{U-O}}(\theta) + (r_{D/P} - r_{D/S}) \times F_{D_{U-O}}(\theta) = 0 \tag{10}$$

Substituting (8) into (10) and performing simplifications leads to:

$$M_{S_A}(\theta) = M_{P_{O-U}}(\theta) + r_{P/S} \times F_{P_{O-U}}(\theta) \tag{11}$$

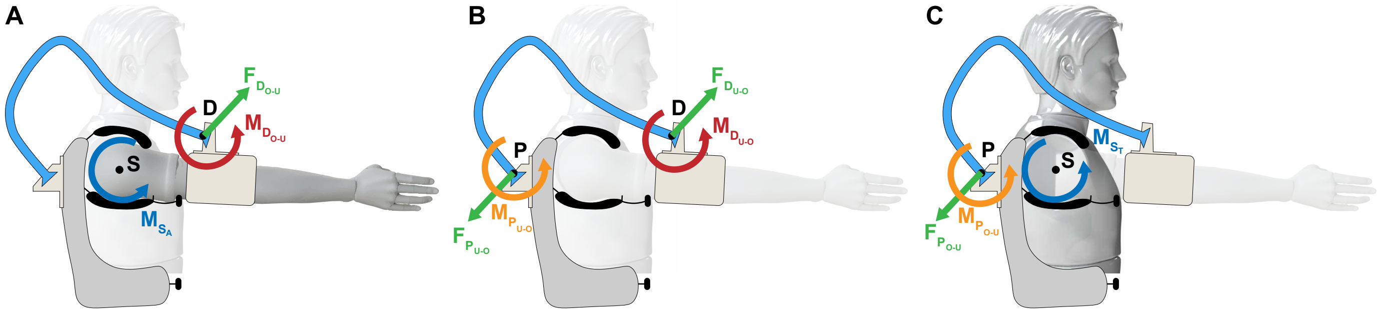


Fig. S5. Free-body schematics detailing the resulting moments and forces experienced on the total system once the orthosis is deformed. A: interaction dynamics between the orthosis and the arm. B: Interaction dynamics solely within the orthosis. C: interaction dynamics between the orthosis and the torso.

Fig. S5C shows a free-body-schematic of the torso relating how the output loading at the proximal end of the orthosis relate to the moment felt at the shoulder from the orthosis. This relationship can be seen below as:

$$M_{S_T} = -M_{P_{O-U}} - r_{P/S} \times F_{P_{O-U}} \tag{12}$$

In which case it is seen that $M_{S_A}$ and $M_{S_T}$ are in fact equivalent quantities with opposite signs, hence demonstrating why converting nodal loads from the proximal end of the orthosis is valid to calculate the compensation moment at the shoulder.

##### *B. Cross-Platform Communication*

A custom VB.NET application was written in Microsoft Visual Studio 2022 to handle automation of the SolidWorks simulation, as well as data transfer to and from the MATLAB optimization. This section will offer a brief description of the tasks performed in this application. All SolidWorks automation was performed using the SolidWorks Application Programming Interface.

The application begins by reading the JavaScript Object Notation file created in MATLAB, containing orthosis contour information and the extrusion parameter ($w$). The contour of the orthosis is automatically sketched, and then extruded to a 3D body using this information. An axis of rotation is placed, corresponding to the shoulder, constrained to cross the point $(X, Y = 0)$, and align normal to the $X - Y$ plane.

The application then automatically creates the nonlinear static study, applies appropriate boundary conditions, and creates a mesh, as described in the primary body of this work. The analysis is then automatically run.

The following information is then written to another JSON, to be interpretted in MATLAB and used to calculate optimization objectives: for each time point - nodal force vectors, nodal displacement vectors, nodal normal, shear, and principle stresses as well as VonMises stress and maximum principle stress intensity, and size of each deformation step.

#### APPENDIX C
#### BENCHTOP VALIDATION

##### A. Benchtop Apparatus

This validation was done on the instrumented benchtop apparatus seen in Fig. S6. The apparatus frame was made from aluminum T-slotted framing rails (McMaster-Carr, Elmhurst, IL, USA). A bearing-shaft housing was 3D printed (Formlabs - Somerville, MA, USA - White Resin), and an Aluminum 6061 rectangular beam was press-fitted onto the shaft. The rotating beam is meant to replicate the load of a scaled-down human arm.

To fix the orthoses to the benchtop apparatus (and similarly to users), dovetails are added to the proximal and distal ends, such that they can be fixed in attachment components with corresponding slots. The trapeziodal shape of the dovetails prevents the orthoses from slipping out of the manufactured clamps due to in-plane deformations. These clamps were screwed into rectangular plates, each of which had an arrays of threaded holes to account for a wide range of attachment points, and were either 3D printed or manufactured (Aluminum 6061). Between the distal attachment and the aluminum beam is an ATI-mini 40 6-axis force-torque transducer (ATI, Novanta Inc., Bedford, MA, USA, sampling frequency 1000 Hz), used to record interaction loading between orthosis and beam.

Small-scale experimental validations were performed to quantify simulation accuracy. Specifically, this was done to validate the overall output compensation moment of the orthoses, validate the individual moment component profiles (e.g., concentrated distal moment), and compare experimental to expected material stiffnesses.

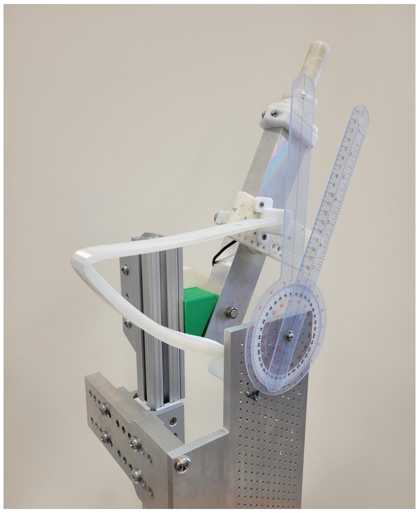

Fig. S6. The small-scale benchtop apparatus used to perform experimental validation. A force transducer is embedded into the setup at the junction of the prototype and the rotating beam.

##### B. Orthosis Fabrication

Several orthosis designs were used for experimental validation. 2 are presented here: Specimen 1 was outputted from an optimization with equivalent settings to Stage 2 presented in our primary work; Specimen 2 was a direct output of Stage 3. These designs were scaled down to a size capable of being printed in most commercially available 3D printers, and re-simulated. These orthoses were then printed on either Ultimaker S5 or Argo 500 printers, using TPU 95A filament (Ultimaker S5) or Flex TPU (Argo 500) for Specimens 1 and 2 respectively.

##### C. Experimental Protocol

A 3D printed handle (Fig. S6) was attached to the end of the aluminum beam. Starting with the beam raised up vertically, with the orthosis attached, the beam was rotated by the handle by increments of 30° until the beam was completely inverted ($\theta = 180°$), upon which the sequence was repeated in reverse. Each position was statically held, and force-torque transducer readings were captured for at least 3 seconds at each pose. A goniometer was fixed to the apparatus such that it aligned with the axis of rotation to provide references for each rotational increment.

##### D. Data Analysis

F/T transducer recordings were filtered using a zero-phase digital low-pass first order Butterworth filter with a cutoff frequency of 20 Hz. Time-series data for each static hold was averaged.

As the transducer lies between the orthosis and aluminum beam, when the beam is rotated and held by the experimenter at the handle, and the orthosis is deformed, transducer recordings are solely due to orthosis reaction. For a given static position, the mean orthosis compensation moment can be calculated as

$$M_G(\theta) = M_{S'}(\theta) + r \times F_{S'}(\theta) \tag{13}$$

that is, the sum of the in-plane moment recorded by the transducer $M_{S'}$, as well as the moment at the axis of rotation caused by the resultant in-plane force vector measured at the transducer $F_{S'}$, where $r$ is the position vector of the transducer relative to the axis of rotation.

As the F/T transducer is not necessarily aligned with the distal orthosis end, the concentrated moment at the distal end is not equivalent to $M_{S'}$. This is because when not aligned, in-plane moments recorded at the transducer due to the orthosis can be due to both the distal moment as well as the result force at the distal end. To determine the distal moment $M_D$, the portion of $M_{S'}$ due to the resultant force $F_{S'}$ is first determined as:

$$M_{S'_{res}}(\theta) = r' \times F_{S'}(\theta) \tag{14}$$

where $r'$ is the position vector of the transducer relative to the distal end of the orthosis. $M_D$ can then be solved as:

$$M_D(\theta) = M_{S'}(\theta) - M_{S'_{res}}(\theta) \tag{15}$$

##### *E. Results*

As scaling differences could be attributed to material and manufacturing processes, the goal of this validation was to confirm the shapes of both the compensation moment profiles $M_G(\theta)$ and the distal moment profiles $M_D(\theta)$. As is seen in Fig. S7, experimental trends follow those simulated; that is, compensation moments (black) are roughly sinusoid, and distal moments (red) follow the simulated trends of increasing over the rotation. Relative proportions of distal moment to compensation moment are also similar between experimental and simulated moments.

Experimental results suggest that Specimen 1 was less stiff than expected, and Specimen 2 was more stiff than expected. As such, to develop a stiffness ratio between experimental and simulated compensation moments, towards identifying if material properties should be altered for improved accuracy within simulation, the slope between moments was estimated using linear regression. A slope of approximately 1 would indicate simulation and experimental results largely agree, while a slope greater than 1 would indicate experimental behavior is stiffer than simulated. Results from this analysis were used to inform future simulations.

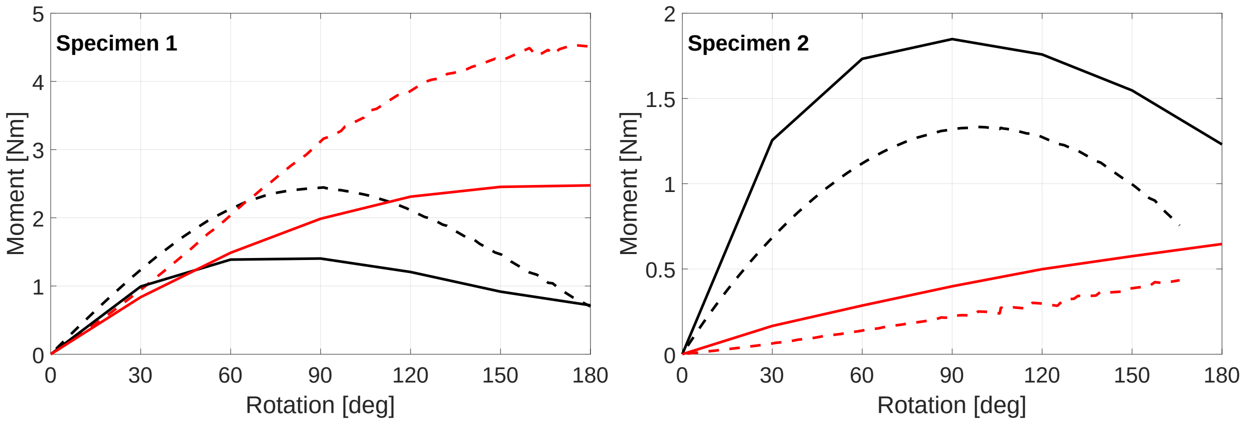


Fig. S7. Benchtop validation results of two orthosis prototypes. Solid lines represent experimental results, while dashed lines represent those simulated. Black lines correspond to overall compensation moment, while red lines correspond to concentrated moments at the orthosis distal ends.

### APPENDIX D
### OPTIMIZATION

#### A. Particle Swarm Optimization

Particle swarm optimization is a genetic algorithm useful for evaluating problems with high-dimensional design vectors [21]. For a single objective function, particle swarm optimization begins by initializing $p$ particles, each corresponding to individual design vectors. At every iteration, the objective function is evaluated based on each particle. These evaluations affect how the positions of the particles will be updated in parameter space for the subsequent iteration. The position update vector $V$ of each particle is based on three components: a particle's own inertia (current trajectory), a cognitive component (a particle's best evaluation over all iterations), and a social component (the cumulative best evaluation over all particles and iterations). This update can be seen below:

$$V_i^{t+1} = c_1 V_i^t + c_2 r_1 (P_i^t - X_i^t) + c_3 r_2 (G^t - X_i^t) \tag{16}$$

where $i$ and $t$ represent the current particle and iteration respectively, $X$ is particle position, $P$ is the particle position at particle $i$'s best evaluation, and $G$ is the particle position for the overall best evaluation. $c_1$, $c_2$, and $c_3$ are weighting constants used to place emphasis on specific components of the update function; $r_1$ and $r_2$ are random coefficients between 0 and 1 updated every iteration [21]. For the implementation used in this work, $c_2$ and $c_3$ are both set to default values of 1.49. $c_1$ is an adaptive constant that modulates based on the rate of new global leaders being determined Specifically, a counter variable ($Count_{inertia}$, initial value of 0) decreases by 1 each iteration a new global leader is selected, and increases by 1 each iteration no new leader is found. If $Count_{inertia} < 2$, $c_1$ is doubled, while if $Count_{inertia} > 5$, $c_1$ is halved. The default $c_1$ range of [0.1,1.1] is used.

#### B. Nonlinear Constraints

*1) Undeformed Intersections:* While the orthosis encoding process ensures the top and bottom edges do not self-intersect (knots are solved), this constraint ensures that the two edges do not intersect with each other.

*2) Deformed Intersections:* This constraint is evaluated based on the deformed shape of the orthosis, returned to the optimizer following simulation. For this evaluation, the edges are checked for intersections with one another and with themselves in the final deformed state of the orthosis.

*3) Incomplete Rotations:* In certain cases, the simulator cannot solve for further displacement steps after a certain amount of deformation. Examples of these cases occur at buckling or limit points, where displacements grow large under constant forces. In such cases, it is assumed that rotating the orthosis further will lead to yielding or other part failures. To increase the number of feasible solutions for considerations, all solutions that have completed 165° or greater are retained.

*4) Initial Collision:* This constraint is used to evaluate if the orthosis can lie in the same plane as the user's arm. Evaluation is performed by defining a polygon to be an estimate of the user's arm. If any point defining the orthosis's contour lies within the arm polygon, the solution is penalized.

*5) Shoulder Collision:* This constraint is used to evaluate if the orthosis can lie in the same plane as the user throughout deformation. Once simulation has completed, the deformed orthosis contour is compared to a polgyon used as an estimate of the user's shoulder. If any point of the orthosis contour lies within the polygon, the solution is penalized.

*6) Final Collision:* Similarly to shoulder collisions, this constraint is used to ensure the orthosis does not collide with the arm after deformation. Once simulation has completed, the deformed orthosis contour is compared to a polgyon used as an estimate of the user's arm at the final rotation. If any point of the orthosis contour lies within the polygon, the solution is penalized.

*7) Exceeded Printer Bounds:* When approaching fabrication steps, it becomes increasingly important to ensure orthosis solutions identified by the optimizer can in fact be easily manufactured. As such, a nonlinear constraint evaluating if the orthosis contour can fit inside commercially available 3D printers was used to ensure all evaluated solutions could be fabricated. The minimum bounding rectangle of the orthosis contour is determined. The side lengths of this rectangle are compared to the X and Y dimensions of a 3D printer's build volume (for example, the Argo 500 by Roboze, with X and Y dimensions of 0.5 x 0.5 m). If rectangle dimensions exceed printer constraints, the solution is penalized.

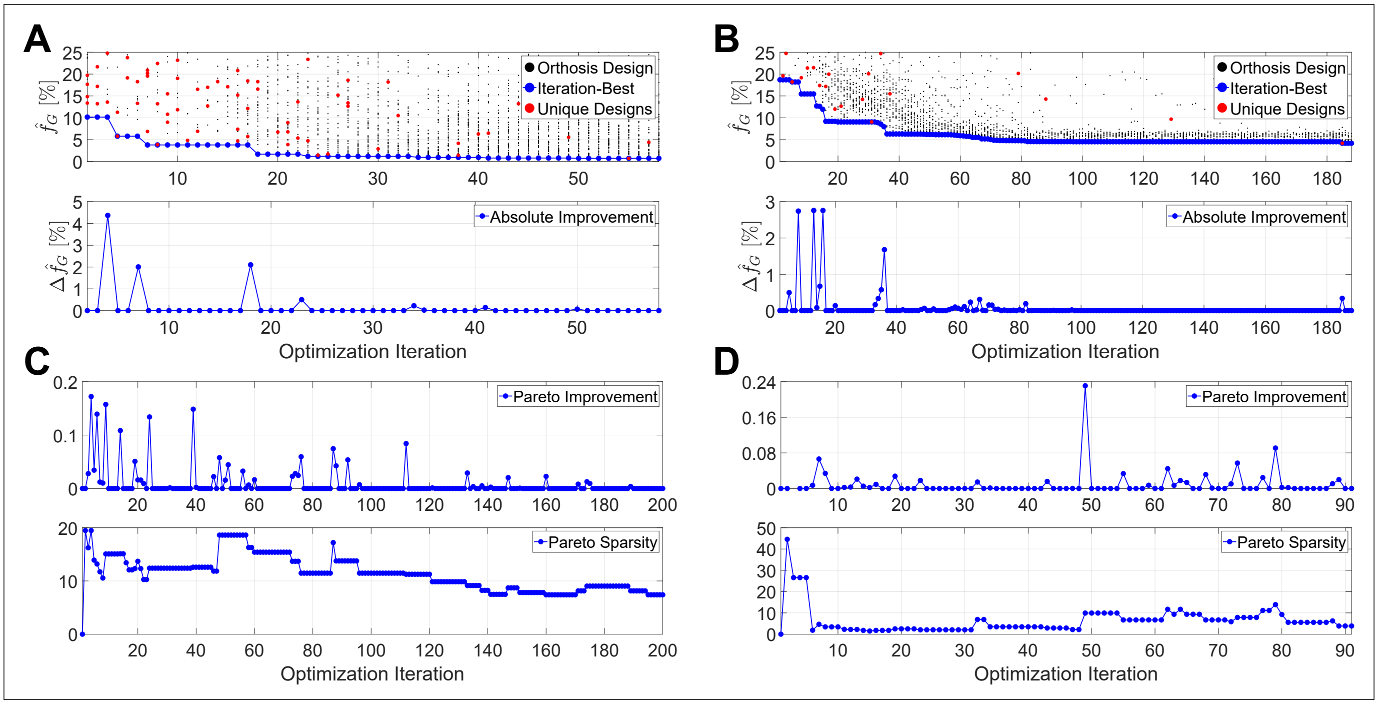


Fig. S8. A & B: Stages 1 and 2 optimization results and convergence respectively. A & B Top: optimization solutions per iteration, with iteration-best solutions highlighted. A & B Bottom: absolute improvement over iteration. C & D: Stages 3 and 4 optimization convergence metrics respectively. C & D Top: Pareto improvement represents the average movement of the Pareto front caused by new Pareto solutions at each iteration. C & D Bottom: Pareto sparsity represents the average distance between neighboring solutions on the Pareto front.

##### *C. Optimizer Convergence*

Fig. S8 show convergence for optimization Stages 1-4. For single-objective optimizations, convergence is defined based on absolute improvement, as in the difference between the optimizer best solution at iteration $i$ and iteration $i-1$.

For multi-objective optimization, convergence was based on improvement and sparsity. Improvement was defined as the following: at each iteration $i$ where a new Pareto solution is found, the minimum distance between each new Pareto solution and the Pareto front at iteration $i-1$ was determined, and averaged across all new Pareto solutions. As such, improvement was defined as the average distance between new Pareto solutions at iteration $i$ and the Pareto front at iteration $i-1$. Sparsity was defined as the mean distance between each neighboring Pareto solutions at each iteration. Both improvement and sparsity are unitless quantities.

##### *D. Optimization Stage 5: Multi-Objective (3), In-Plane, dimensional*

*1) Methods:* As a preliminary exploration into the extension of this framework to higher dimensions, a final optimization was run using three objectives: $\hat{f}_{G_{Dim}}$, $\hat{f}_M$, and a third: Orthosis Protrusion $\hat{f}_P$. Optimization settings, constraints, and design parameter bounds are the same as those used in Stage 4.

The same set of 11 solutions used in the inital generation of the Stage 4 optimization were used in the initialization for this optimization, with the remaining 39 particles being randomly generated.

*a) Orthosis Protrusion:* Towards limiting the size of the orthoses to increase wearability, a size objective was defined based on the maximum perpendicular distance of the orthosis from the user's midline. This was done by calculating the perpendicular distance of each point on the undeformed orthosis's contour to the user's midline, and extracting the maximum as:

$$\hat{f}_P = \frac{\max\limits_{i=1,\ldots,n} P_i}{0.6} \tag{17}$$

where $\hat{f}_P$ represents the normalized maximum protrusion of the orthosis, $n$ represents the number of points used to define the orthosis contour, and 0.6 is used for normalization, roughly corresponding to the 50$^{\text{th}}$ percentile male arm length [22].

*2) Results:* This optimization was run for 231 iterations, evaluating 11,550 orthosis designs. Fig. S9 shows the results of this optimization as a comparison with Stage 4. As is seen, the solution improves in $\hat{f}_P$ relative to Stage 4 (where $\hat{f}_P$ was not optimized for); while solutions can score similarly in $\hat{f}_{G_{Dim}}$, solutions with similar $\hat{f}_M$ are not found. Two designs are highlighted for presentation: Design G ($\hat{f}_{G_{Dim}} = 5.0\%$, $\hat{f}_M = 49.5\%$, $\hat{f}_P = 85.6\%$) and Design H ($\hat{f}_{G_{Dim}} = 25.7\%$, $\hat{f}_M =$

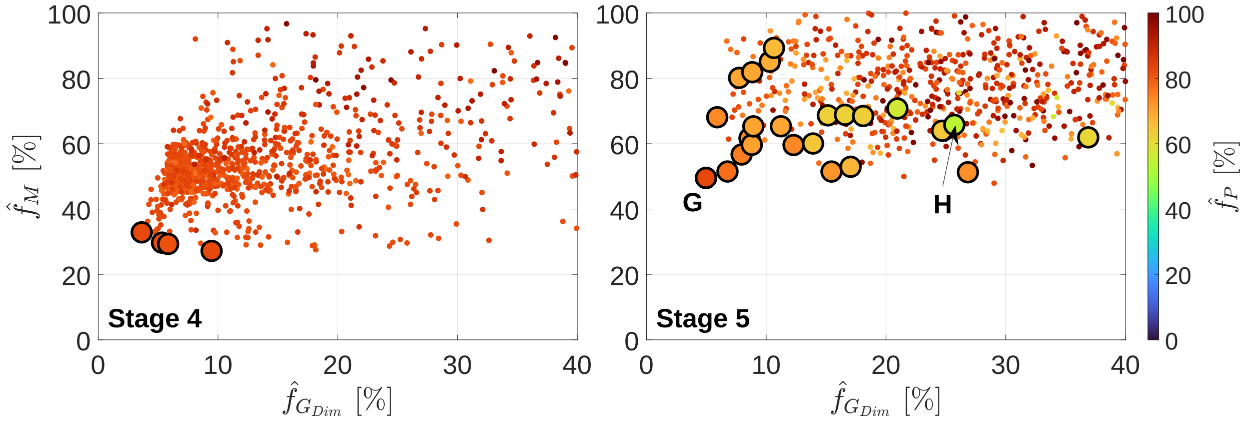


Fig. S9. Optimization Stages 4 (left) and 5 (right) comparison. Pareto solutions are seen as represented by their objective scores. While Stage 5 is the only optimization stage to incorporate all three objectives, Stage 4 is presented here with $\hat{f}_P$ scores colored for comparison.

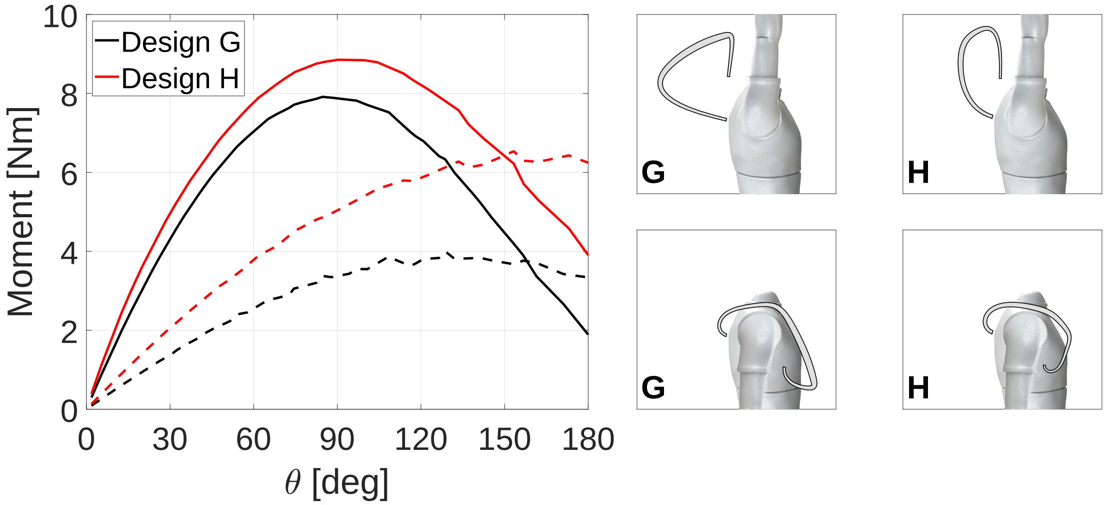


Fig. S10. Stage 5 optimization results. Left: moment profiles for Designs G (black) and H (red). Solid lines represent overall compensation moment. Dashed lines represent concentrated distal moment. The undeformed and fully deformed configurations of Designs G and H are seen in the middle and right columns respectively.

$65.8\%$, $\hat{f}_P = 53.8\%$). Design G was selected as, within the constrained range of objective scores, it scored the best in $\hat{f}_{G_{Dim}}$ and $\hat{f}_M$; Design H was selected as the design with the best $\hat{f}_P$ (Fig. S10).

### APPENDIX E
### ORTHOSIS FABRICATION

The Argo 500 3D printer by Roboze was used for manufacturing, due to its larger than standard build volume (500 x 500 x 500 mm). Flex TPU filament was selected for manufacturing, based on its suitable contrast between stiffness and elongation prior to failure (Young's modulus and Poisson's ratio as listed in the primary work, Ultimate tensile strength in-plane: 28 MPa, elongation at ultimate tensile strength: 400 %). To account for printer size and material, the following scaling was applied to Design E to target a moment amplitude of about 8 Nm: the entire 2D profile of the orthosis, including attachment points, was scaled by 0.75$\times$; in-plane thickness was scaled by 0.79$\times$; out-of-plane width was scaled up to 4 cm. The resulting amount of compensation provided by this orthosis was estimated to be 8.46 Nm.

Design F was selected from the output of Stage 4 (multi-objective, in-plane, dimensional). This orthosis was also printed with Flex TPU on the Argo 500. As this optimization was dimensional, and included printer constraints, no post-design scaling was performed. The resulting amount of compensation provided by this orthosis was estimated to be 8.07 Nm.

#### A. Orthosis-User Interface

The orthosis-user interface was designed such that the orthosis can be worn on the right arm. The orthosis-user interface can be thought of as two junctions: the proximal and distal. The proximal attachment is a modified backpack-based harness made by IUVO (Pontedera PI, Italy) for the Active Pelvis Orthosis. This harness was selected as it secures to the user's chest, shoulders, and waist, therefore offloading applied loads at many points. The harness also features a rigid plate on the back, making it possible to build off and attach the orthosis. This was done using 3D printed components (BambuLabs - Austin, TX, USA - PLA, Formlabs - Somerville, MA, USA - White Resin) that were attached onto the harness's rigid plates.

The arm junction consists of an adjustable arm cuff, with an orthosis attachment slot fixed on top. The arm cuff was made by collaborators at Independence Prosthetics and Orthotics (Newark, DE, USA) and is primarily made out of rigid plastic, with padding underneath. Velcro straps are used for adjustment.

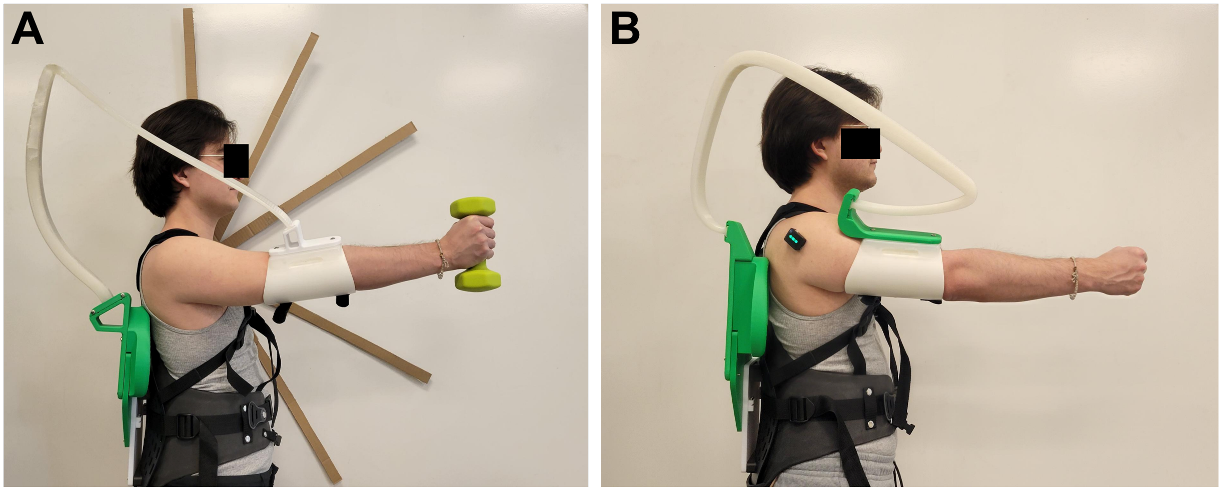


Fig. S11. A: One of this article's authors wearing the orthosis Design E, holding the weight used in experiments in front of the reference targets. B: One of this article's authors wearing the orthosis Design F.

### APPENDIX F
### IN-VIVO TESTING

#### A. *Muscle Activity - Orthosis Design E*

Fig. S12 shows the group average results from Static trials for the PM, UT, and PD from orthosis Design E. Annotations indicate the results of the four-way Static mixed model. Fig. S11 showcases orthosis Design E, with the participant holding the weight used for the experiment in front of the reference targets.

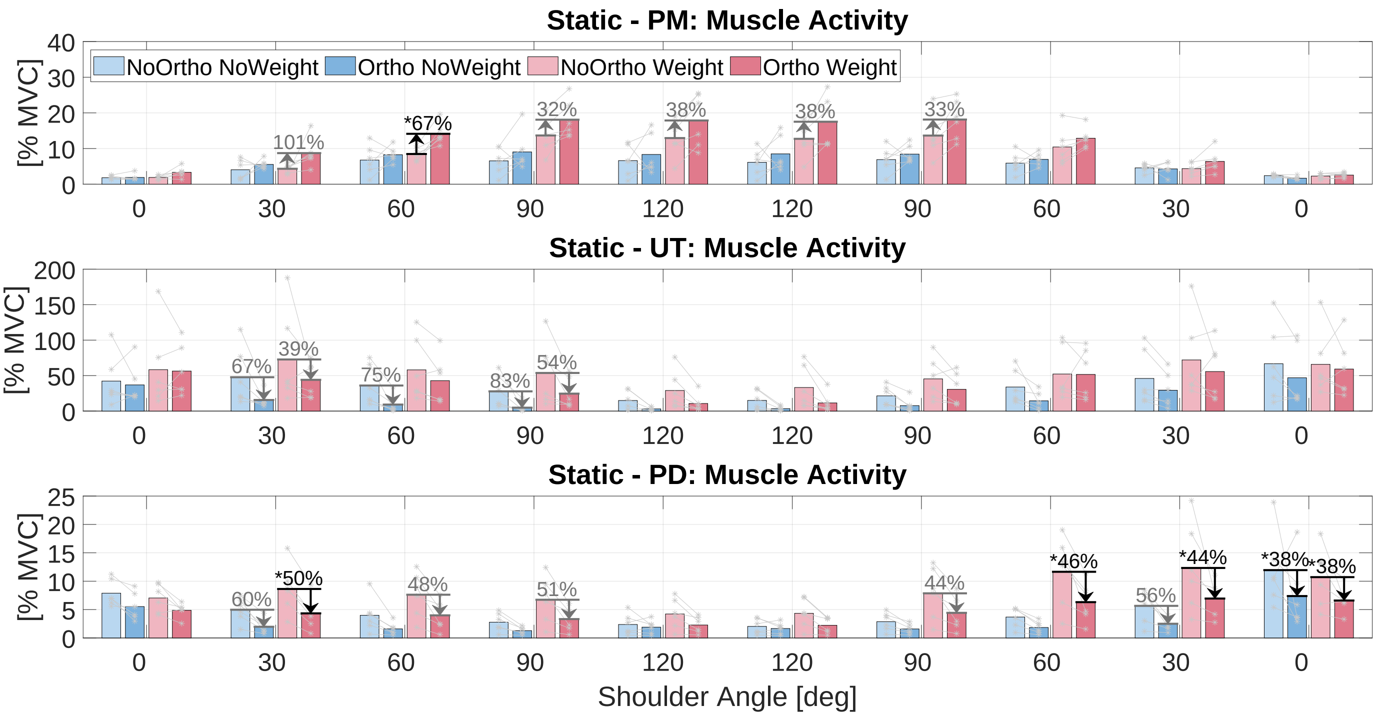


Fig. S12. Muscle activity measured during static tasks using orthosis Design E. From top to bottom: breakdown of muscle activity by shoulder angle and posture repetition for the PM, UT, and PD in order. Individual participants are notated by asterisks, with lines joining repeated participants. Black arrows indicate significant changes in muscle activity due to wearing the orthosis after Bonferroni correction.

*1) Linear Mixed Models:* Results of the four-way linear mixed model for static trials, the three-way linear mixed model for static trials, and the four-way linear mixed model for dynamic trials are seen in Tables S2, S3, and S4 respectively.

TABLE S2
LINEAR MIXED MODEL RESULTS FOR STATIC TRIALS - ALL POSTURES AND REPETITIONS

| Effect | F Stat | $DF_{Num}$ | $DF_{Den}$ | p Value |
|---|---|---|---|---|
| **AD Muscle Activity** | | | | |
| Orthosis | 40.068 | 1 | 195 | <**0.0001** |
| Weight | 151.201 | 1 | 195 | <**0.0001** |
| Orthosis*Weight | 7.049 | 1 | 195 | **0.0086** |
| Posture | 44.383 | 4 | 195 | <**0.0001** |
| Orthosis*Posture | 3.078 | 4 | 195 | **0.0173** |
| Weight*Posture | 9.082 | 4 | 195 | <**0.0001** |
| Orthosis*Weight*Posture | 0.335 | 4 | 195 | 0.8539 |
| Rep | 10.266 | 1 | 195 | **0.0016** |
| Orthosis*Rep | 4.098 | 1 | 195 | **0.0443** |
| Weight*Rep | 0.142 | 1 | 195 | 0.7063 |
| Orthosis*Weight*Rep | 0.849 | 1 | 195 | 0.3580 |
| Posture*Rep | 2.117 | 4 | 195 | 0.0801 |
| Orthosis*Posture*Rep | 0.694 | 4 | 195 | 0.5968 |
| Weight*Posture*Rep | 1.050 | 4 | 195 | 0.3827 |
| Orthosis*Weight*Posture*Rep | 0.494 | 4 | 195 | 0.7399 |
| **PM Muscle Activity** | | | | |
| Orthosis | 34.607 | 1 | 195 | <**0.0001** |
| Weight | 131.014 | 1 | 195 | <**0.0001** |
| Orthosis*Weight | 9.091 | 1 | 195 | **0.0029** |
| Posture | 88.215 | 4 | 195 | <**0.0001** |
| Orthosis*Posture | 2.159 | 4 | 195 | 0.0750 |
| Weight*Posture | 16.313 | 4 | 195 | <**0.0001** |
| Orthosis*Weight*Posture | 0.149 | 4 | 195 | 0.9631 |
| Rep | 0.504 | 1 | 195 | 0.4784 |
| Orthosis*Rep | 1.676 | 1 | 195 | 0.1970 |
| Weight*Rep | 0.006 | 1 | 195 | 0.9407 |
| Orthosis*Weight*Rep | 0.194 | 1 | 195 | 0.6598 |
| Posture*Rep | 0.116 | 4 | 195 | 0.9767 |
| Orthosis*Posture*Rep | 0.303 | 4 | 195 | 0.8757 |
| Weight*Posture*Rep | 0.233 | 4 | 195 | 0.9198 |
| Orthosis*Weight*Posture*Rep | 0.150 | 4 | 195 | 0.9628 |
| **UT Muscle Activity** | | | | |
| Orthosis | 42.854 | 1 | 195 | <**0.0001** |
| Weight | 63.024 | 1 | 195 | <**0.0001** |
| Orthosis*Weight | 0.285 | 1 | 195 | 0.5942 |
| Posture | 30.383 | 4 | 195 | <**0.0001** |
| Orthosis*Posture | 0.986 | 4 | 195 | 0.4165 |
| Weight*Posture | 1.884 | 4 | 195 | 0.1147 |
| Orthosis*Weight*Posture | 0.676 | 4 | 195 | 0.6093 |
| Rep | 2.090 | 1 | 195 | 0.1499 |
| Orthosis*Rep | 1.007 | 1 | 195 | 0.3168 |
| Weight*Rep | 0.150 | 1 | 195 | 0.6994 |
| Orthosis*Weight*Rep | 0.090 | 1 | 195 | 0.7641 |
| Posture*Rep | 0.746 | 4 | 195 | 0.5614 |
| Orthosis*Posture*Rep | 0.800 | 4 | 195 | 0.5266 |
| Weight*Posture*Rep | 0.254 | 4 | 195 | 0.9069 |
| Orthosis*Weight*Posture*Rep | 0.072 | 4 | 195 | 0.9906 |
| **PD Muscle Activity** | | | | |
| Orthosis | 89.148 | 1 | 195 | <**0.0001** |
| Weight | 71.873 | 1 | 195 | <**0.0001** |
| Orthosis*Weight | 6.161 | 1 | 195 | **0.0139** |
| Posture | 33.850 | 4 | 195 | <**0.0001** |
| Orthosis*Posture | 2.484 | 4 | 195 | **0.0451** |
| Weight*Posture | 11.886 | 4 | 195 | <**0.0001** |
| Orthosis*Weight*Posture | 0.613 | 4 | 195 | 0.6535 |
| Rep | 20.527 | 1 | 195 | <**0.0001** |
| Orthosis*Rep | 1.159 | 1 | 195 | 0.2829 |
| Weight*Rep | 5.021 | 1 | 195 | **0.0262** |
| Orthosis*Weight*Rep | 0.329 | 1 | 195 | 0.5671 |
| Posture*Rep | 2.869 | 4 | 195 | **0.0243** |
| Orthosis*Posture*Rep | 0.410 | 4 | 195 | 0.8010 |
| Weight*Posture*Rep | 1.206 | 4 | 195 | 0.3096 |
| Orthosis*Weight*Posture*Rep | 0.132 | 4 | 195 | 0.9704 |

TABLE S3
LINEAR MIXED MODEL RESULTS FOR STATIC TRIALS - GROUPED ROM

| Effect | F Stat | $DF_{Num}$ | $DF_{Den}$ | p Value |
|---|---|---|---|---|
| **AD Muscle Activity** | | | | |
| Orthosis | 18.786 | 1 | 227 | <**0.0001** |
| Weight | 73.461 | 1 | 227 | <**0.0001** |
| Orthosis*Weight | 4.323 | 1 | 227 | **0.0387** |
| PostureType | 23.329 | 1 | 227 | <**0.0001** |
| Orthosis*PostureType | 3.487 | 1 | 227 | 0.0631 |
| Weight*PostureType | 8.244 | 1 | 227 | **0.0045** |
| Orthosis*Weight*PostureType | 0.273 | 1 | 227 | 0.6016 |
| **PM Muscle Activity** | | | | |
| Orthosis | 24.303 | 1 | 227 | <**0.0001** |
| Weight | 83.508 | 1 | 227 | <**0.0001** |
| Orthosis*Weight | 7.174 | 1 | 227 | **0.0079** |
| PostureType | 271.878 | 1 | 227 | <**0.0001** |
| Orthosis*PostureType | 5.678 | 1 | 227 | **0.0180** |
| Weight*PostureType | 47.670 | 1 | 227 | <**0.0001** |
| Orthosis*Weight*PostureType | 0.214 | 1 | 227 | 0.6438 |
| **UT Muscle Activity** | | | | |
| Orthosis | 36.452 | 1 | 227 | <**0.0001** |
| Weight | 53.416 | 1 | 227 | <**0.0001** |
| Orthosis*Weight | 0.323 | 1 | 227 | 0.5704 |
| PostureType | 79.876 | 1 | 227 | <**0.0001** |
| Orthosis*PostureType | 0.051 | 1 | 227 | 0.8214 |
| Weight*PostureType | 0.115 | 1 | 227 | 0.7344 |
| Orthosis*Weight*PostureType | 0.131 | 1 | 227 | 0.7176 |
| **PD Muscle Activity** | | | | |
| Orthosis | 64.310 | 1 | 227 | <**0.0001** |
| Weight | 42.235 | 1 | 227 | <**0.0001** |
| Orthosis*Weight | 3.390 | 1 | 227 | 0.0669 |
| PostureType | 64.435 | 1 | 227 | <**0.0001** |
| Orthosis*PostureType | 3.185 | 1 | 227 | 0.0756 |
| Weight*PostureType | 3.633 | 1 | 227 | 0.0579 |
| Orthosis*Weight*PostureType | 0.749 | 1 | 227 | 0.3877 |

TABLE S4
LINEAR MIXED MODEL RESULTS FOR DYNAMIC MOVEMENTS

| Effect | F Stat | $DF_{Num}$ | $DF_{Den}$ | p Value |
|---|---|---|---|---|
| **AD Muscle Activity** | | | | |
| Orthosis | 16.557 | 1 | 75 | <**0.0001** |
| Weight | 138.182 | 1 | 75 | <**0.0001** |
| Orthosis*Weight | 1.987 | 1 | 75 | 0.1628 |
| Trial | 79.312 | 1 | 75 | <**0.0001** |
| Orthosis*Trial | 1.288 | 1 | 75 | 0.2601 |
| Weight*Trial | 8.714 | 1 | 75 | **0.0042** |
| Orthosis*Weight*Trial | 0.569 | 1 | 75 | 0.4531 |
| Speed | 6.059 | 1 | 75 | **0.0161** |
| Orthosis*Speed | 0.056 | 1 | 75 | 0.8128 |
| Weight*Speed | 0.499 | 1 | 75 | 0.4821 |
| Orthosis*Weight*Speed | 0.153 | 1 | 75 | 0.6970 |
| Trial*Speed | 3.744 | 1 | 75 | 0.0568 |
| Orthosis*Trial*Speed | 0.002 | 1 | 75 | 0.9675 |
| Weight*Trial*Speed | 0.058 | 1 | 75 | 0.8107 |
| Orthosis*Weight*Trial*Speed | 0.000 | 1 | 75 | 0.9985 |
| **PM Muscle Activity** | | | | |
| Orthosis | 23.335 | 1 | 75 | <**0.0001** |
| Weight | 168.252 | 1 | 75 | <**0.0001** |
| Orthosis*Weight | 0.112 | 1 | 75 | 0.7384 |
| Trial | 44.058 | 1 | 75 | <**0.0001** |
| Orthosis*Trial | 0.312 | 1 | 75 | 0.5779 |
| Weight*Trial | 12.596 | 1 | 75 | **0.0007** |
| Orthosis*Weight*Trial | 0.000 | 1 | 75 | 0.9991 |
| Speed | 3.150 | 1 | 75 | 0.0800 |
| Orthosis*Speed | 1.128 | 1 | 75 | 0.2915 |
| Weight*Speed | 0.042 | 1 | 75 | 0.8391 |
| Orthosis*Weight*Speed | 0.401 | 1 | 75 | 0.5285 |
| Trial*Speed | 0.587 | 1 | 75 | 0.4459 |
| Orthosis*Trial*Speed | 0.168 | 1 | 75 | 0.6828 |
| Weight*Trial*Speed | 0.240 | 1 | 75 | 0.6256 |
| Orthosis*Weight*Trial*Speed | 0.008 | 1 | 75 | 0.9311 |
| **UT Muscle Activity** | | | | |
| Orthosis | 21.243 | 1 | 75 | <**0.0001** |
| Weight | 57.051 | 1 | 75 | <**0.0001** |
| Orthosis*Weight | 0.037 | 1 | 75 | 0.8484 |
| Trial | 25.014 | 1 | 75 | <**0.0001** |
| Orthosis*Trial | 2.005 | 1 | 75 | 0.1609 |
| Weight*Trial | 4.582 | 1 | 75 | **0.0356** |
| Orthosis*Weight*Trial | 0.068 | 1 | 75 | 0.7955 |
| Speed | 0.882 | 1 | 75 | 0.3507 |
| Orthosis*Speed | 0.088 | 1 | 75 | 0.7671 |
| Weight*Speed | 0.197 | 1 | 75 | 0.6587 |
| Orthosis*Weight*Speed | 0.054 | 1 | 75 | 0.8172 |
| Trial*Speed | 0.662 | 1 | 75 | 0.4184 |
| Orthosis*Trial*Speed | 0.049 | 1 | 75 | 0.8248 |
| Weight*Trial*Speed | 0.007 | 1 | 75 | 0.9344 |
| Orthosis*Weight*Trial*Speed | 0.008 | 1 | 75 | 0.9274 |
| **PD Muscle Activity** | | | | |
| Orthosis | 38.062 | 1 | 75 | <**0.0001** |
| Weight | 127.595 | 1 | 75 | <**0.0001** |
| Orthosis*Weight | 2.305 | 1 | 75 | 0.1332 |
| Trial | 26.202 | 1 | 75 | <**0.0001** |
| Orthosis*Trial | 1.981 | 1 | 75 | 0.1635 |
| Weight*Trial | 5.509 | 1 | 75 | **0.0216** |
| Orthosis*Weight*Trial | 0.007 | 1 | 75 | 0.9341 |
| Speed | 1.462 | 1 | 75 | 0.2305 |
| Orthosis*Speed | 0.028 | 1 | 75 | 0.8674 |
| Weight*Speed | 0.029 | 1 | 75 | 0.8656 |
| Orthosis*Weight*Speed | 0.010 | 1 | 75 | 0.9221 |
| Trial*Speed | 1.402 | 1 | 75 | 0.2402 |
| Orthosis*Trial*Speed | 0.032 | 1 | 75 | 0.8592 |
| Weight*Trial*Speed | 0.414 | 1 | 75 | 0.5217 |
| Orthosis*Weight*Trial*Speed | 0.009 | 1 | 75 | 0.9238 |

#### *B. Muscle Activity - Orthosis Design F*

Figs. S13, S14, and S15, show participant-specific results for the participant that repeated the experiment using orthosis Design F. Fig S11 shows the participant wearing orthosis Design F.

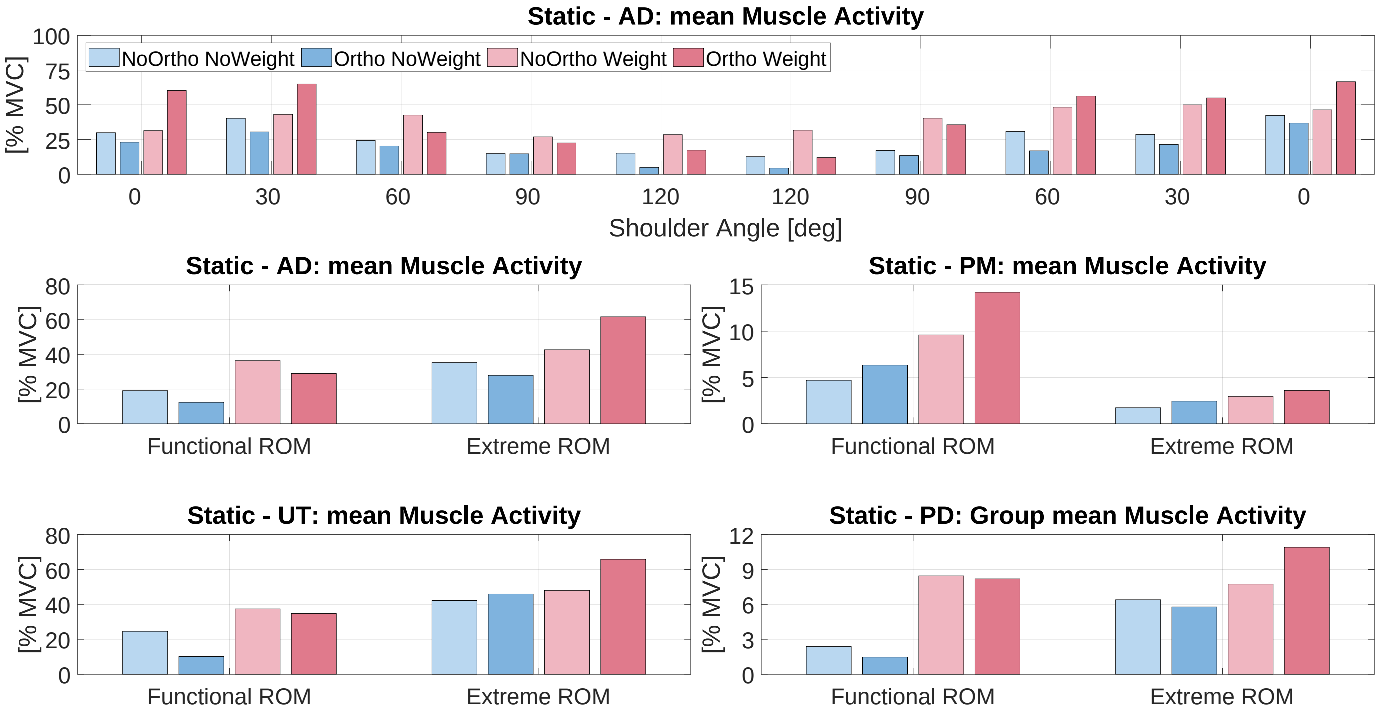


Fig. S13. Muscle activity measured during static tasks by a single participant using the orthosis Design F. Top row: breakdown of AD muscle activity by shoulder angle and posture repetition. The second and third rows show muscle activity for all muscles, grouped by posture type: Functional vs. Extreme.

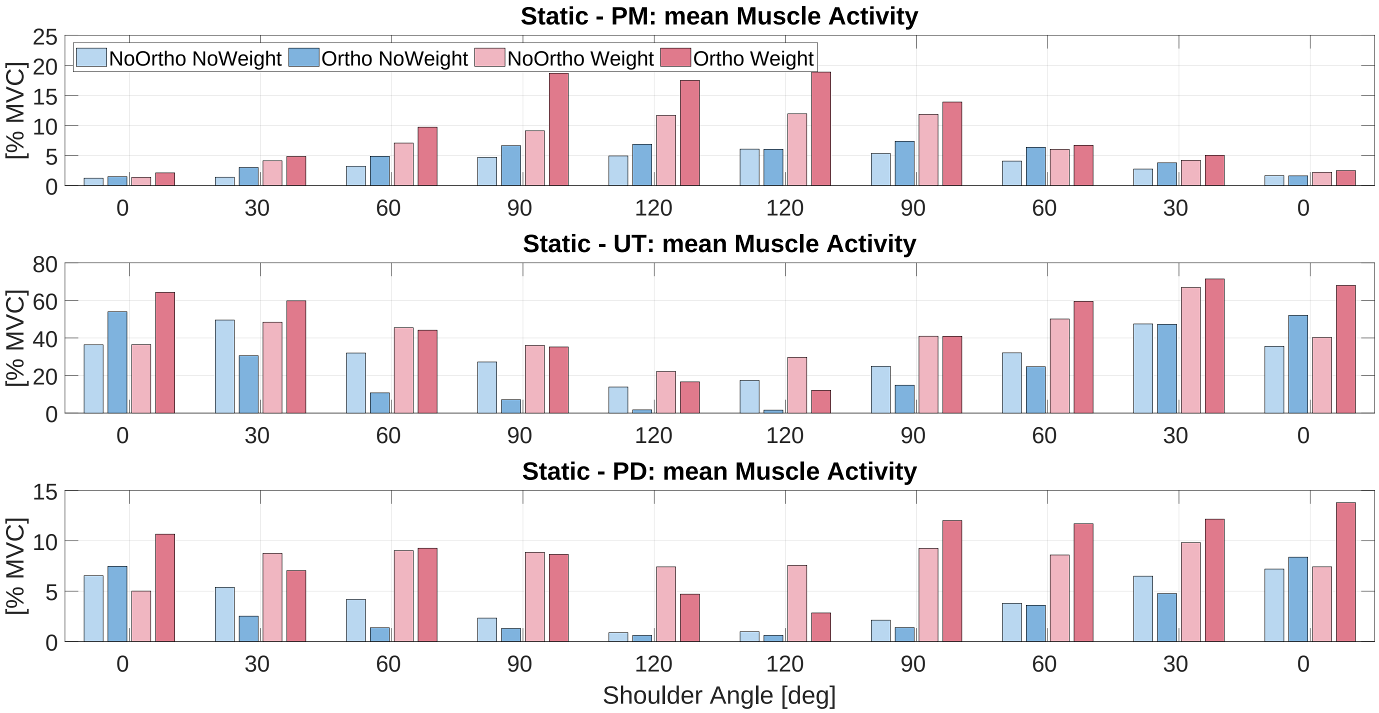


Fig. S14. Muscle activity measured during static tasks by a single participant using the orthosis Design F. From top to bottom: breakdown of muscle activity by shoulder angle and posture repetition for the PM, UT, and PD in order.

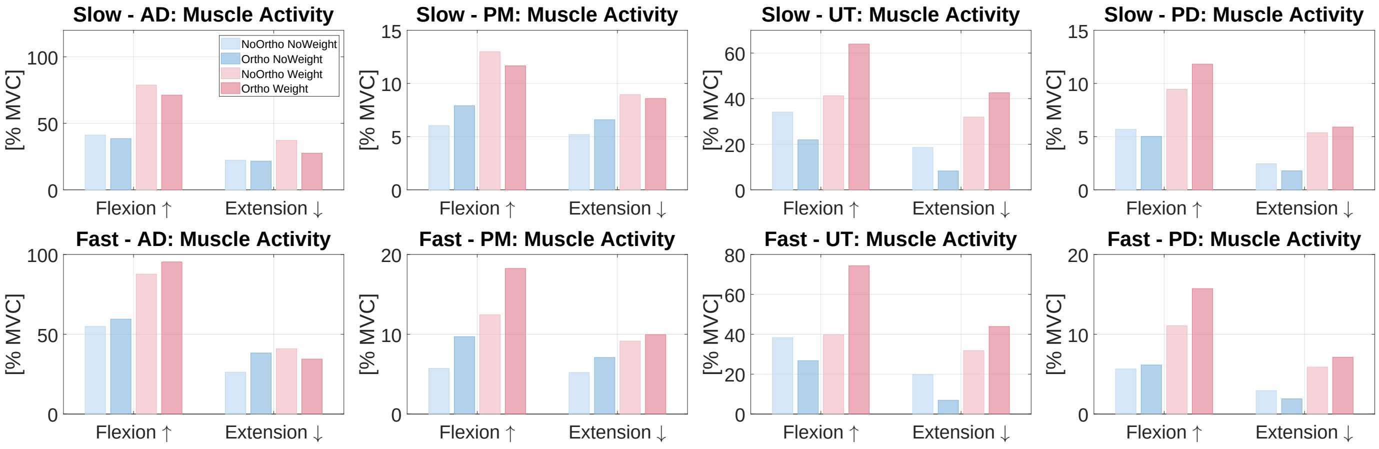


Fig. S15. Muscle activity measured during dynamic tasks by a single participant using the orthosis Design F. Results are plotted by condition combination, movement type, and speed. The columns from left to right show results for AD, PM, UT, and PD in order. Top row shows results for Slow movements, while the bottom row shows results for Fast movements.

#### C. *NASA-TLX*

The specific questions posed as part of the modified NASA-TLX questionnaire are enumerated below. Participant responses for each domain are seen in Table S5, alongside group average responses.

1) Physical demand: how much physical activity was required? (e.g., was the task easy or demanding, slow or brisk, slack or strenuous, restful or laborious)
2) Performance: how successful do you think you were in accomplishing the goals of the task?
3) Effort: how hard did you have to work (mentally and physically) to accomplish your level of performance?
4) Frustration: how frustrated did you feel during the task? (e.g., insecure, discouraged, irritated, stressed, and annoyed versus secure, gratified, content, relaxed, and complacent)

TABLE S5
NASA TLX SURVEY RESULTS

| Design E | P01 | P02 | P03 | P04 | P05 | P06 | Average |
|---|---|---|---|---|---|---|---|
| Physical | 3 | 4 | 4 | 3 | 4 | 2 | 3.3 |
| Performance | 1 | 2 | 2 | 4 | 1 | 3 | 2.2 |
| Effort | 2 | 4 | 2 | 4 | 1 | 3 | 2.7 |
| Frustration | 1 | 1 | 3 | 1 | 1 | 1 | 1.3 |

Participant P01, who repeated the experiment with Design F, had the following responses for the second experiment: Physical - 5, Performance - 2, Effort - 4, Frustration - 2.
